# Rapgef1 paralog-mediated regulation of Wnt/β-catenin signaling orchestrates early embryo tissue patterning and morphogenesis

**DOI:** 10.1101/2024.11.21.624777

**Authors:** Tuhina Prasad, Sharada Iyer, Sian D’silva, Reuben J. Mathew, Divya Tej Sowpati, Vegesna Radha, Megha Kumar

## Abstract

RAPGEF1, a guanine nucleotide exchange factor, regulates signaling and cytoskeletal dynamics in mammalian cells, yet its role in development remains unclear as *Rapgef1* null mouse embryos do not survive beyond implantation. We demonstrate that zebrafish rapgef1 is maternally expressed, and its paralogs, rapgef1a and rapgef1b, exhibit tissue and developmental stage-specific splicing. Disruption of *rapgef1b* caused brain and somite defects, impaired cranial neural crest specification, and microcephaly-like phenotypes, uncovering its previously uncharacterized functions in morphogenesis and tissue patterning. Transcriptomic analyses and differential gene expression provide fresh insights into the developmental functions of *rapgef1b* in presomitic mesoderm and somitogenesis by modulating the Wnt/β catenin signaling. Rapgef1b deficient embryos also showed spindle pole disorganization and chromosome mis-congression, linking Rapgef1 to centrosome-mediated mitotic fidelity. Together, our findings identify Rapgef1b as a key regulator of neural crest development, mesodermal morphogenesis, and early mitoses, highlighting its tissue-specific functions during vertebrate embryogenesis.

**Teaser:** In this study, we show that *rapgef1b* is essential for shaping the embryonic brain, somites, and body axis by regulating gene expression, cell division, survival, and differentiation. Our findings reveal a new dimension of signaling-mediated cell fate specification during early vertebrate development.

## Introduction

Guanine Nucleotide Exchange Factors (GEFs) serve as key signaling centers that activate small GTPases, enabling multiple effector functions to regulate adhesion, cytoskeletal dynamics, and signaling. One of the GEFs, RAPGEF1 (C3G), is predominantly expressed as a 140kD protein with a modular architecture consisting of a C-terminal catalytic domain, a central protein interaction region enriched in proline-rich Crk binding motifs (CBR), and an N-terminal E-cadherin binding domain (*1*). RAPGEF1 localizes to the subcortical cytoskeleton, Golgi, nuclear speckles, centrosome, and exhibits nucleo-cytoplasmic exchange (*2–5*). Its activity is tightly controlled through intramolecular interactions, post-translational modifications such as phosphorylation, and intermolecular protein interactions. RAPGEF1 functions as an enzyme and a scaffold by engaging with diverse signaling proteins (*3*, *6*). At the molecular level, it interacts with GRB2, CRK, CAS, Src family kinases, β-catenin, GSK3β, and cenexin in mammalian cells, regulating chromatin organization, splicing, cytoskeletal remodeling, and transcription (*2*, *5*, *7*, *8*). Deregulated *RAPGEF1* levels and missense mutations are associated with multiple human pathologies, including cancers and neurological disorders (*9–17*). Although tissue-specific, alternately spliced isoforms of *Rapgef1* have been described in mice, their functions remain unknown (*18*, *19*). Notably, a novel brain-specific, long isoform of *Rapgef1* has recently been identified in adult murine brains and human brain organoids (*7*, *18–21*).

The loss-of-function studies underscore the developmental importance of *Rapgef1*. Mouse embryonic stem cells lacking *Rapgef1* showed enhanced self-renewal but failed to undergo lineage differentiation (*22*, *23*). Further, *Rapgef1* knockout mice embryos failed to survive beyond implantation, highlighting its essential role in early embryogenesis (*24*, *25*). Hypomorphic mice expressing a human *Rapgef1* allele exhibited severe defects in cell adhesion, vasculogenesis, and differentiation (*26–28*). However, the role of *rapgef1* in early embryonic events such as cell fate specification, lineage determination, tissue patterning and morphogenesis remains unresolved.

To address this gap, we investigated *rapgef1* function in zebrafish, which possess two paralogs, *rapgef1a* and *rapgef1b*. Knockdown of *rapgef1a* has been reported to cause pan developmental defects, including brain and blood vessel anomalies, motor neuron defects, and impaired locomotor activity (*13*). In contrast, the function of the other homolog, *rapgef1b,* has not been explored. Here, we examined the spatiotemporal expression of transcripts generated from the two paralogs and demonstrate that *rapgef1a* and *rapgef1b* are both expressed in early embryogenesis, showing differences in isoform-specific expression patterns during development and in adult tissues. We delineate an essential role of *rapgef1b* in regulating spindle pole integrity and architecture coupled with faithful chromosome segregation during early mitoses. Further, our results highlight that *rapgef1b* is indispensable for embryo survival, maintaining progenitor fate in cranial neural crest cells and head size. Remarkably, *rapgef1b* also aids in orchestrating presomitic mesoderm patterning and somitogenesis via modulation of Wnt/ β-catenin signaling. This study positions *rapgef1b* as a pivotal regulator of early embryogenesis, integrating β-catenin -mediated lineage specification of paraxial mesoderm and its derivatives.

## Results

### *Rapgef1* is expressed during embryonic development and in adult tissues of zebrafish

Zebrafish paralogs, *rapgef1a,* and *rapgef1b*, located on chromosomes 8 and 21 respectively, show high homology in the coding regions and encode proteins with similar domain organization (Fig. 1A). Both paralogues contain the highly conserved Crk binding region (CBR) which includes a hotspot for additional exons (orange arrowhead, Fig. 1A). The expression of *rapgef1* during embryonic development and in adult tissues was examined using primers corresponding to the sequence in the 5’ end (N-ter), for the amplification of *rapgef1a* and *1b* specific gene products (Fig. 1B, S1A). These primers are common to multiple isoforms and are predicted to amplify products of 130bp and 167bp from *rapgef1a* and *1b,* respectively. Semi-quantitative PCR resulted in amplicons of expected sizes from all the developmental stages (32-cell to 2 days post fertilization, dpf). Our results show that transcripts from both paralogs are expressed during early developmental stages, such as the 32-cell stage, indicating that rapgef1 mRNA is maternally deposited. The expression of *rapgef1* continues into the mid-blastula stage (256 cell-3.3hpf stage) and somitogenesis phase (6 - 21somite stage) until 2dpf (Fig. 1B). A recent report has shown that depletion of Rapgef1a results in gross developmental defects, brain and vasculature abnormalities (*13*). However, the role of other paralog, Rapgef1b remains unknown. Hence, we focused on investigating the developmental functions of this paralog Rapgef1b in early embryogenesis. First, to delineate the spatiotemporal and developmental gene expression profile of *rapgef1b,* we used *in situ* hybridization (ISH) with sequence-specific antisense riboprobes, which revealed its ubiquitous expression across developmental stages (red asterisk, Fig 1C), with higher anterior expression at 1dpf (red arrowhead, Fig. 1C). Specificity of the *rapgef1b* expression was confirmed by using sense riboprobes as control (Fig. 1C). Western blotting showed the expression of a 150kD polypeptide of Rapgef1, similar to that seen in mammalian cells and tissues, in all the early developmental stages (Fig. 1D). Brain tissue from adult zebrafish expressed a distinct polypeptide, indicating tissue-specific expression of alternate isoforms as described earlier using mouse tissues and alluding to isoform specific role in neural functions (Fig. S1B) (*18*).

**Figure 1:**
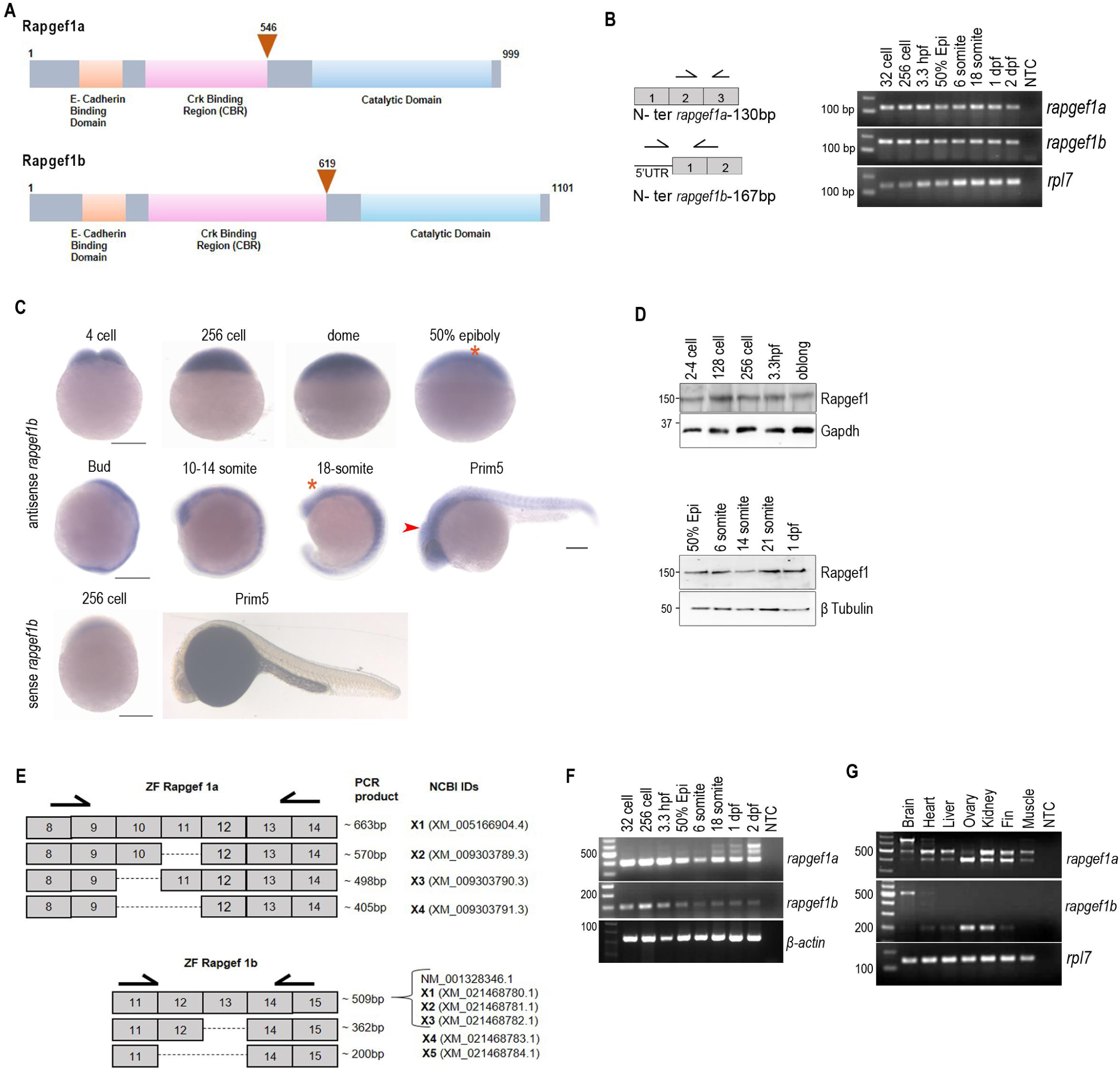
Expression profile of *rapgef1* and its isoforms during embryogenesis and adult zebrafish tissues. A: Schematic illustration showing the protein domains of zRAPGEF1a and zRAPGEF1b, triangle (orange) indicates hotspot for additional exons in the CBR region. Numbers indicate total amino acids and the position of insertion of additional amino acids in the alternate isoforms. B: Schematic representation of *rapgef1a* and *1b* 5’ end with the primer positions (black arrows) and the exon numbers. RT-PCR analysis of *rapgef1a* and *1b* transcripts in different developmental stages. Zebrafish *rpl7* (*ribosomal protein L 7*) was the loading control. NTC represents no template control. C: Whole-mount RNA *in situ* hybridization showing *rapgef1b* expression during early development. Scale bars, 200µm. D: Western blot showing Rapgef1 levels during embryonic development. β-Tubulin and Gapdh were used as the loading controls. E: Schematic representation of *rapgef1a* and *b* exons showing predicted alternatively spliced transcripts for *rapgef1a* and *1b* with nucleotide ID (https://www.ncbi.nlm.nih.gov/nuccore). Locations of isoform-specific primers (black arrows) are shown that give products of different lengths based on the presence or absence of additional exons in alternately spliced isoforms. F: RT-PCR analysis of *rapgef1* transcripts in different developmental stages using *rapgef1a* and *1b* isoform-specific primers. β actin was used as the loading control. G: Isoform-specific amplification of *rapgef1a* and *1b* transcripts in adult tissues (brain, heart, liver, ovary, fin, muscle). Zebrafish *rpl7* was used as the loading control. NTC represents no template control.

### Differential isoform-specific expression of *rapgef1* paralogs in embryonic and adult tissues

Expression of a higher molecular weight isoform of Rapgef1 in the adult zebrafish brain (Fig. S1B) indicates generation of alternate isoforms, including a set of cassette exons, may be expressed. Zebrafish *rapgef1* transcripts span 3257bp (*rapgef1a*), and 3566bp (*rapgef1b*), with the longest variants are composed of 25 (*rapgef1a*) and 28 (*rapgef1b*) exons. Publicly available databases like NCBI suggest the generation of alternatively spliced isoforms through the inclusion of a pair of exons (10 & 11 in *rapgef1a* and 12 & 13 in *rapgef1b*). Isoform-specific primers for both *rapgef1a* and *1b* which are expected to generate PCR products of different sizes based on the inclusion of one or both exons, were used to study the expression of alternate isoforms (Fig.1E). *Rapgef1a* transcript lacking the additional exons (X4, 405bp product) is expressed throughout embryonic development till early somitogenesis (6 somite stage) (Fig.1F). Interestingly, expression of the longer isoforms of *rapgef1a* with 1 or more exons, (X3-498bp, X2-570bp, X1-633bp amplicons) begins with late somitogenesis (18-21 somite stage) (Fig.1F). The longest isoform with exons 10 & 11 is predominantly expressed in the juvenile head and adult brain, with poor expression of other products suggesting the isoform-specific function of *rapgef1a* in the adult brain (Fig. S1C). Other adult tissues, such as the ovary, heart, liver, and kidney show expression of various isoforms in addition to the embryonic isoform (Fig.1G).

In contrast, only the smallest isoform lacking exons 12 and 13 (X5, 200bp amplicon) of *rapgef1b* is expressed until 2dpf and its levels decrease as the embryo develops (Fig. 1F). Remarkably, there is dynamic isoform switching which results in the expression of the brain-specific, longer isoform of *rapgef1b* containing exons 12 & 13 in the juvenile and adult brain with age (Fig. S1C). Adult tissues such as the heart, liver, ovary, and kidney predominantly express the isoform lacking cassette exons, with no detectable levels of the longer transcripts of *rapgef1b* (Fig.1G). We also investigated the expression patterns of the brain-specific isoforms by ISH with sequence-specific antisense riboprobe (Fig. S1D). *Rapgef1b* (antisense riboprobe to exon 3 and 4) is expressed in the diencephalon (double red asterisk), rods, and cones (red arrow), ganglion cell layer (red arrowhead), gut, and myotomes of 3dpf larvae (red asterisk) (Fig. S1D). In contrast, the brain-specific isoform expression is restricted to diencephalon, and the ganglion cell layer of the eye, but is absent in rods and cones and myotome (red arrow, Fig.S1D). Thus, the expression of the brain-specific isoform of *rapgef1b* is restricted to the cranial region and absent from mesodermal-derived tissues like the myotome.

### *Rapgef1b* depletion results in neurodevelopmental and anteroposterior axis patterning defects

To determine the functional role of *rapgef1b* in embryogenesis, we used two loss of function strategies, the knockdown approach using sequence-specific antisense morpholino (ATG/ translational blocker) and a CRISPR-based knockout approach against z*Rapgef1b*. The morpholino approach was used for rapid and effective knockdown of gene expression to understand developmental functions during embryogenesis. Further, as morpholino-based knockdown strategy may induce off target effects, we used the CRISPR-based gene editing method and generated crispants (F0) to corroborate the morphant phenotypes (Fig. 2). The *rapgef1b* morphants injected with ATG blocker morpholino and *rapgef1b* crispants showed depletion of Rapgef1 protein. The *rapgef1b* KO fish showed a deletion of 366bp, confirming effective abrogation of its expression (Fig. 2C, S2A). Both the strategies resulted in similar phenotypes, the morphants were characterized as P0 (normal, similar to WT), P1 (brain and axis defects), and P2 (severe brain and axis deformities) (Fig. 2B, 2D, S2D). *Rapgef1b* depletion by both strategies resulted in defects in the brain, eye, and inner ear in 24-hour post-fertilization embryos. The P2 embryos appear smaller in size, showed poorly developed eyes, defects in the brain architecture, and lack a proper midbrain-hindbrain boundary (red asterisk, Fig. 2B). The crispants and morphants also showed somitic defects (red arrowhead, Fig.2B) and anterior-posterior body axis abnormalities, marked by shortening of the axis and tail (red arrow, Fig. 2B). We have used time matched, severe phenotype embryos classified as P2 to perform the comparative gene expression analyses in this study. The severity of the phenotype and the survival rate were observed in a morpholino concentration-dependent manner with time (Fig. S2B, S2C). To ascertain that the phenotype was specific to Rapgef1b knockdown, we performed rescue experiments by introducing human *rapgef1* mRNA, which resulted in partial, but significant rescue (39%) of the severe phenotype, P2 (Fig. 2D, E). In contrast, human *rapgef1* mRNA of a predicted deleterious mutant Y485C (Mutagenetix & ClinVar), a residue conserved in zRapgef1, that shows altered stability and compromised phosphorylation by Abl (unpublished findings from our lab), could not rescue the *rapgef1b*-specific phenotype (Fig. 2D, E). These results suggest that *rapgef1b* is essential for early development and morphogenesis of the brain, eye, somites, as well as the elongation of the body axis and tail.

**Figure 2:**
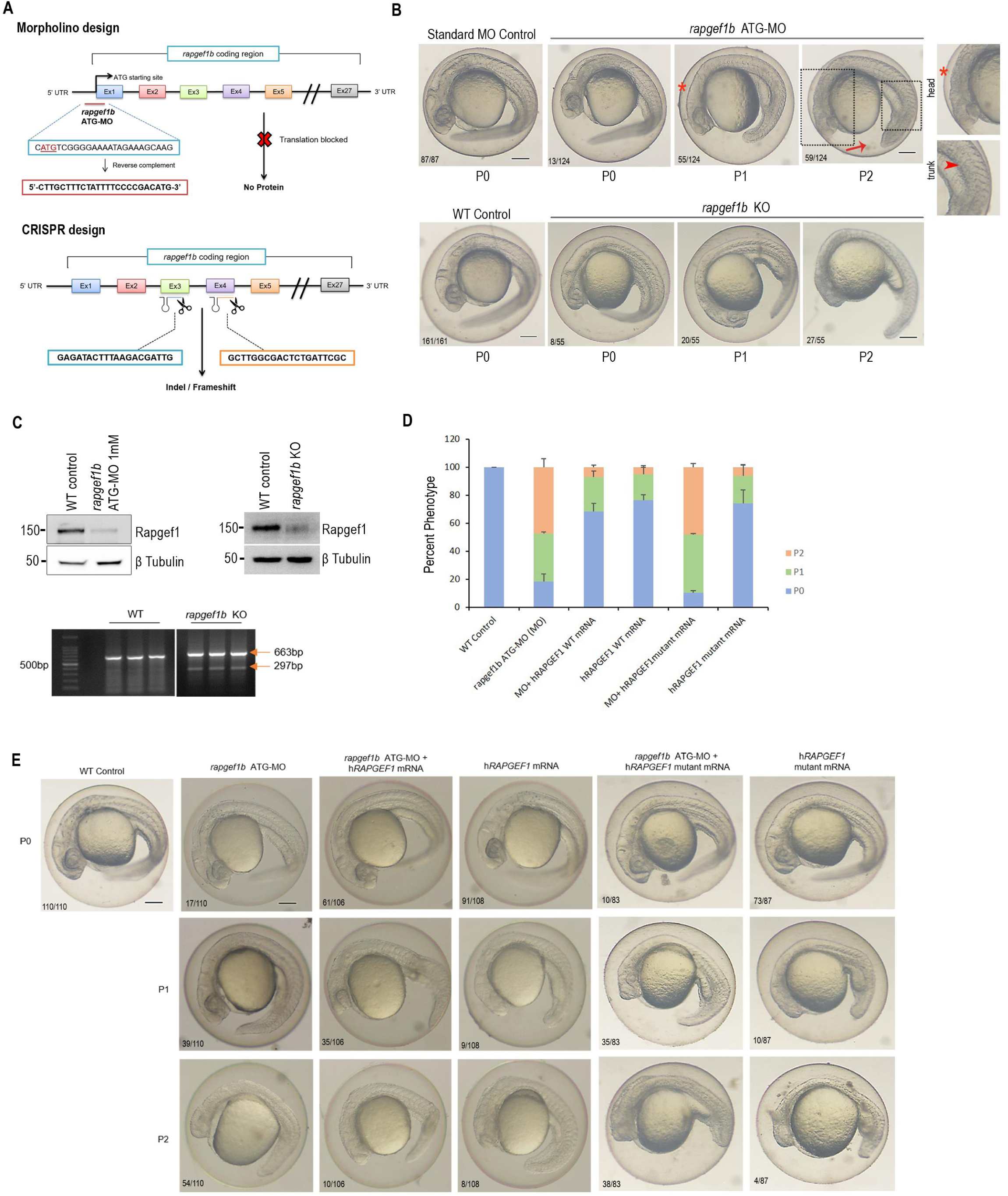
*Rapgef1b* is required for brain and anteroposterior axis development. A: Schematic representation of *rapgef1b* knockdown and knockout strategies by morpholino and CRISPR-Cas9-based approaches respectively. The morpholino binds to the translation start site, abrogating protein synthesis. sgRNA was designed to generate a deletion of 366bp between exons 3 and 4. B: Representative images showing gross morphological appearance of *rapgef1b* depleted embryos by morpholino and CRISPR strategies in comparison to control embryos at 1dpf showing midbrain-hindbrain boundary (red asterisk) and body axis defects (red arrow). The boxed areas show the magnified images of defects in midbrain-hindbrain defects (red asterisk) and body axis defects (red arrow). Scale bars, 500µm. C: Upper panel showing Rapgef1 levels by western blot in *rapgef1b* morpholino injected morphants and *rapgef1b* crispants compared to WT control. β-Tubulin was used as the loading control. The lower panel shows PCR-based analysis to validate the exonic deletion in CRISPR mutants. D: Quantification showing percent phenotype of morphants in rescue experiment. Data are shown as mean ± SEM, n=3 biological repeat experiments, with a minimum of 60 embryos analyzed for each experiment. E: Rescue experiments showing the functional equivalence of human Rapgef1 as a *rapgef1b* ortholog in zebrafish embryos.

### *Rapgef1b* is critical for the maintenance of progenitor fate in cranial neural crest, formation of cranial neural crest derivatives, and maintenance of head size

As *rapgef1b* is expressed in early development and its depletion resulted in brain and anteroposterior axis abnormalities, we inferred that *rapgef1b* is required for early neurodevelopment and axis specification. To test this hypothesis, we performed gene expression analysis of key cranial neural crest specification markers and patterning on *rapgef1b* morphants and crispants. The morphants/crispants showed a significant reduction in in the expression of the neural crest specifier, *twist1a* (red asterisk, Fig. 3A) and the neural plate marker *sox2* (red arrowhead, Fig. 3D). *Twist1a* is expressed in the cranial neural crest-derived head mesenchyme, diencephalon, and is required for the migration of neural crest cells and their differentiation (*29*). *Sox2* is required for otic and epibranchial placode induction, promotes the maintenance of pluripotency, and regulates the development of the sensory epithelium and neural tube (*30*, *31*). The reduced expression of *twist1a* and *sox2* strongly indicates that *rapgef1b* plays a key role in the maintenance of progenitor population in the cranial neural crest, sensory placodes, and is involved in neural tube patterning. We also observed a reduced expression of premigratory neural crest marker, *sox10* (red arrow, Fig. 3B) (*32*, *33*). The expression was rescued in the morphants injected with human *Rapgef1* mRNA (red arrowhead, Fig. 3B). These results suggest that pre-migratory neural crest cells marked by *sox10* expression are reduced in the morphants and may be the underlying cause of a poorly developed brain (Fig.2B). We also observed increased cell death in the anterior region of the embryo (forebrain, midbrain-hindbrain boundary/ MHB, and midbrain) as shown by acridine orange staining (white asterisk, Fig. S3A).

**Figure 3:**
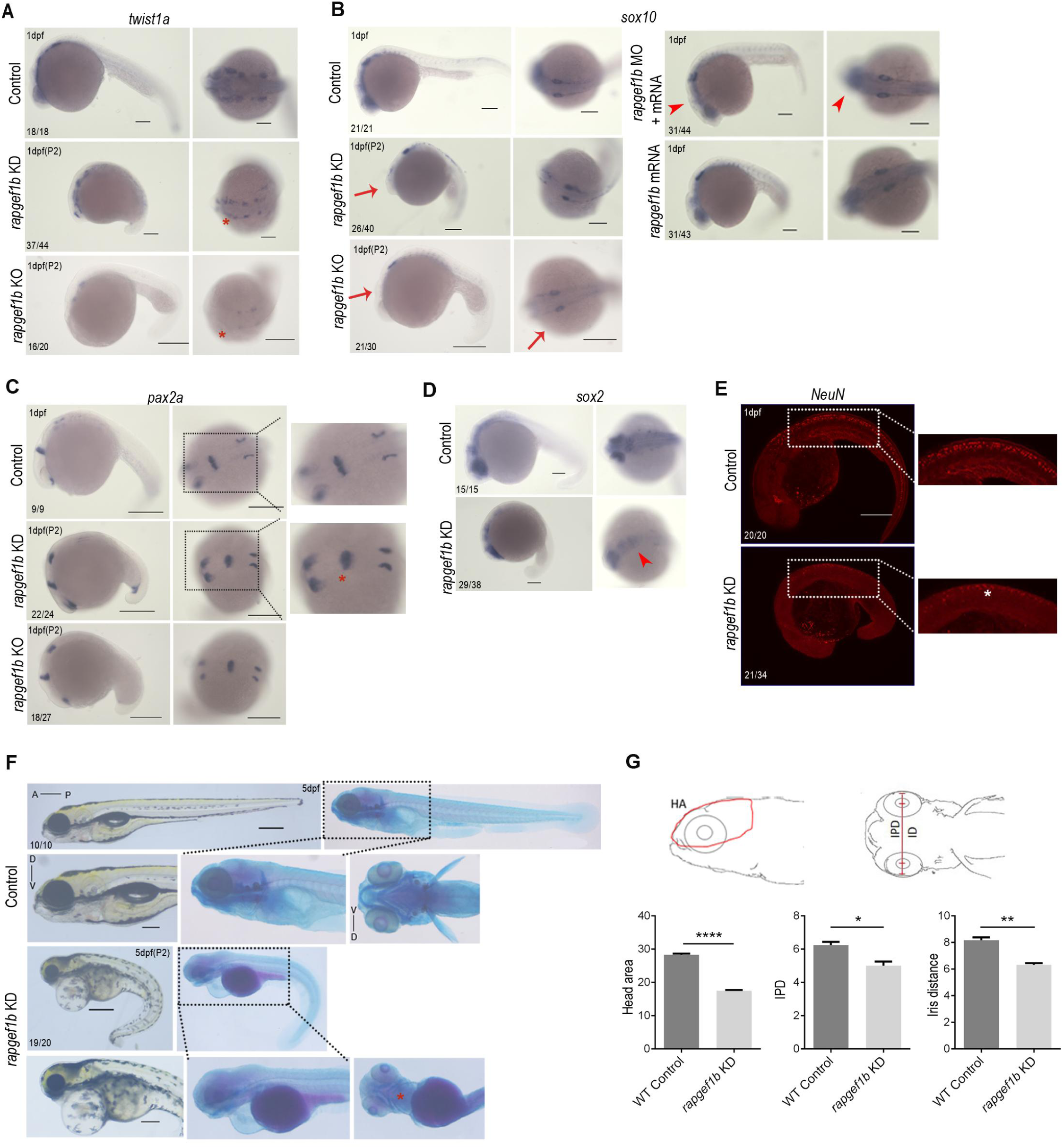
*Rapgef1b* is essential for cranial neural crest specification and maintenance of head size. Gene expression analysis by *in situ* hybridization in control and *rapgef1b* morphants/ mutants at 1dpf. A: *Twist1a* expression (red asterisk) in control and *rapgef1b* morphants/ mutants. B: Expression of *sox10* in *rapgef1b* morphants, mutants (red arrow), and rescue embryos (red arrowhead). Scale bars, 200µm. C: *Pax2a* expression in *rapgef1b* morphants/ mutants showing expression in the optic vesicle, midbrain-hindbrain boundary (red asterisk, inset), and otic vesicle (inset). D: Expression of *sox2* in the neural tube of *rapgef1b* morphants (red arrow). Scale bars, 200µm. E: Control and *rapgef1b* morphants showing post-mitotic neuronal marker neuN (white asterisk). Scale bars, 200µm. F: Skeletal preparations of *rapgef1b* morphants (red asterisk, inset) indicating bone and cartilage staining using Alizarin red S and Alcian blue respectively. Scale bars, 500µm. G: Quantification of microcephaly features – head area (HA), interpupillary distance (IPD), and iris distance (ID) in *rapgef1b* morphants. All data are shown as mean ± SEM. *p<0.05, **p<0.01, ***p<0.001. n=3 biological repeat experiments and the sample size is indicated in each image.

Other key regulators of specification, differentiation, and morphogenesis are the *Pax* family members. *Pax2a* is expressed in the eye, MHB, otic placode, and the pronephros (*34–37*). *Pax2a,* along with *pax8* is required to maintain otic cell fate, and high levels of *pax2a* promote otic differentiation (*38*). The *rapgef1b* morphants exhibit an expansion of *pax2a* expression domains in the eye, otic vesicle, and MHB (red asterisk, Fig.3C). Thus, *rapgef1b* is essential to maintain progenitor cell fate, and its depletion results in an increase in differentiation-inducing signals such as *pax2a*. Concomitantly, we also observed an upregulation of *egr2b*, which is a rhombomere 3 and 5 specifier in the segmented hindbrain (Fig. S3B) (*39–42*). We also examined whether *rapgef1b* is essential for neuronal differentiation. Neuronal nuclei (NeuN) mark the post-mitotic neurons as it is distributed in the nuclei of mature CNS neurons. Thus, NeuN marks terminal differentiation and determination of neuronal phenotype (*43*, *44*). The morphants showed a decrease in NeuN-positive cells (white asterisk, Fig.3E), indicating that depletion of *rapgef1b* results in the decrease of post-mitotic mature neurons and strongly indicates its requirement in terminal neuronal maturation and differentiation.

At larval stages of development (4-5dpf), the *rapgef1b* morphants exhibited cardiac edema, severe craniofacial skeletal anomalies and defects in ceratobranchial skeletal elements (red asterisk, Fig.3F). Craniofacial skeletal elements are derived from the *sox10*-positive cranial neural crest, and the mesenchyme is derived from sclerotome and lateral plate mesoderm (*45*). The *rapgef1b*-depleted embryos showed a reduction in *sox10-*positive neural crest cells, which may be the underlying basis of the observed craniofacial defects. Further, the 5dpf larvae also showed microcephaly-associated craniofacial features such as reduced overall head area (HA), interpupillary distance (IPD), and iris distance (ID) (Fig. 3G), suggesting that rapgef1b plays a key role in formation of craniofacial skeleton and maintenance of overall head size in the embryo.

### *Rapgef1b* plays an important role in paraxial mesoderm differentiation

To further explore the developmental functions of *rapgef1b*, we performed transcriptome sequencing analysis to identify differentially expressed genes (DEGs) and altered processes and pathways in the *rapgef1b* morphants contributing to the observed phenotypes. The principal component analysis (PCA) and correlation matrix showed variance and the agreement of the three replicates across conditions (Fig. S4A, B). The volcano map showed the distribution of DEGs - 76 genes were downregulated and 106 genes were upregulated at the transcriptional level with clear patterns of clustering in the *rapgef1b* deficient embryos within the FDR adjusted p-value <= 0.05 (Fig. 4A). The altered DEGs are listed, and their P values are shown in Fig. S4C. We also performed GO pathway analysis to further investigate the functional relationships between the DEGs. Gene ontology analysis showed an enrichment of anterior/posterior pattern specification, somite development, segmentation, and mesodermal morphogenesis genes indicating that *rapgef1b* plays a key role in paraxial mesoderm specification, patterning and differentiation (Fig. 4B). The heat map shows the selected up-regulated and down-regulated genes in these mentioned processes (Fig. 4C). To validate the transcriptome analysis, we assayed transcript levels of some of the up and down-regulated genes in control and *rapgef1b* morphants (Fig. 4D-H). The *rapgef1b* morphants showed increased transcript levels of the up-regulated group of genes –*msgn1, tbx16*, *cdx4*, *tbx6* and *bbc3*. *Msgn1* and *tbx16* induce the differentiation program in the presomitic mesoderm (PSM) cells by reducing the expression of progenitor maintenance markers *fgf8* and *ntl,* and inducing the expression of *tbx6*. Hence, *msgn1* is essential to regulate the progenitor pool population until somitogenesis is complete (*46*, *47*). *Rapgef1b* morphants showed upregulation of *msgn1* and *tbx16* (Fig. 4D, red asterisk, Fig.4E). We observed an overall increase in *tbx6* transcripts in the morphants, particularly in the tailbud region (red arrow, Fig. 4E). *Tbx6* plays important role in the specification of paraxial mesoderm, somite patterning, and myogenesis (*48–50*). Studies show that ectopic *tbx6* expression promotes less mesodermal fate (*51*). This increase in *tbx6* expression levels results in reduced mesodermal fate and may be the underlying cause of the failure of somite segmentation in the tailbud mesoderm. Thus, upregulation of *msgn1* and *tbx16* in *rapgef1b* morphants by transcriptomic analysis shows that progenitor fate is suppressed and an untimely differentiation program is initiated upon *rapgef1b* depletion (Fig. 4D, 4E).

**Figure 4:**
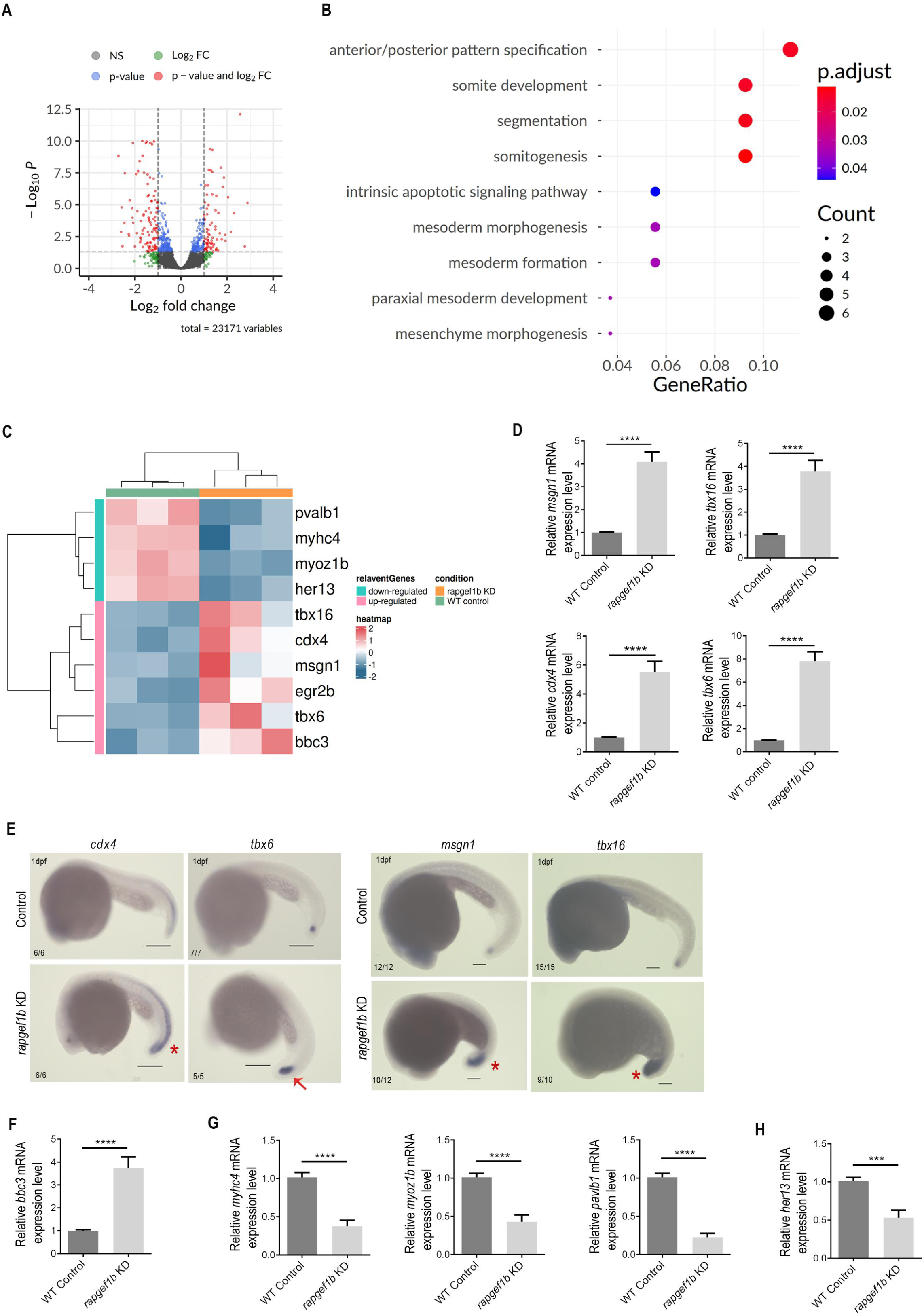
Comparative transcriptome and differential gene expression analysis indicating the role of *rapgef1b* in paraxial mesoderm differentiation. **A:** Volcano plot of the differentially expressed genes (DEGs) in control and *rapgef1b* morphants. Red dots represent DEGs with fold change (>1 or < -1) with p-value <0.05. Green dots represent genes that are significant with fold change (>1 or < -1) with a p-value >0.05. Blue dots represent genes with insignificant fold changes with p-value <0.05. Black dots represent the genes that show insignificant levels of DEGs expression by standards of p-value and log fold change. B: Dot plot representing the gene ontology (GO) analysis of genes upregulated in the *rapgef1b* morphants. C: Heatmap of differentially expressed genes (DEGs) in control and *rapgef1b* morphants. The colors are mapped to z-score normalized values of the log fold change across each gene set. D: Validation by qPCR of *msgn1*, *tbx16, cdx4* and *tbx6* upon *rapgef1b* depletion. Data are shown as mean ± SEM, *p<0.05, **p<0.01, ***p<0.001. n=3 for each biological repeat experiment. E: Gene expression patterns of *cdx4* (red asterisk), *tbx6* (red arrow), msgna1 (red asterisk) and tbx16 (red asterisk) in *rapgef1b* morphants compared to control. Scale bars, 200µm. F: Validation by qPCR of *bbc3* upon *rapgef1b* depletion. G: Validation by qPCR of *myhc4, myoz1b,* and *pavlb1* upon *rapgef1b* depletion. H: Validation by qPCR of *her13* on *rapgef1b* depletion. All data are shown as mean ± SEM. *p<0.05, **p<0.01, ***p<0.001. n=3 for each experiment. Each n represents 100 embryos.

Early mesodermal cell fate is also regulated by the *cdx* family, which also plays key roles in axial elongation, anteroposterior patterning, and the development of the trunk and tail (*52*). Differentiation of paraxial mesoderm and somitogenesis requires other *tbx* family members as well, along with Wnt and FGF signalling (*48*, *51*, *53–55*). *Cdx4* is also an early trunk neural crest specifier, expressed in the trunk neural crest cell progenitors and the developing tailbud (*56*, *57*). Interestingly, *rapgef1b*-depleted embryos show an upregulation of *cdx4* transcript levels in transcriptome analysis, qPCR and *in situ* hybridization (Fig. 4C, 4D, red asterisk, Fig. 4E), suggesting that trunk neural crest cells are maintained in the progenitor fate in the morphants. This led us to conclude that *rapgef1b* is essential to initiate differentiation fate in the trunk neural crest population. Our results show that *rapgef1b* is essential to maintain the progenitor cell fate of the cranial neural crest (Fig.3), however, it is required for the initiation of the differentiation fate in the mesodermal lineage (Fig.4).

We also observed upregulation of intrinsic apoptotic genes such as *bbc3* in our transcriptome analysis and qPCR validation (Fig. 4F) (*58*, *59*). Hence, pro-apoptotic mRNAs are upregulated in the *rapgef1b* morphants, which results in increased apoptotic cells, also marked by acridine orange in the cranial region of *rapgef1b* morphants (Fig. S3A). In contrast, *rapgef1b* morphants showed reduced transcript levels of skeletal muscle genes such as *myhc4*, *myoz1b*, and *pvalb1* (Fig. 4G) (*60–62*). Further, *her13* which is required for the formation of all somitic borders was also downregulated in the morphants (Fig. 4H). The differences in expression of the above genes is likely the underlying basis of disrupted somite architecture in the *rapgef1b* morphants (red arrow, Fig. 2B) (*63*). Hence, transcriptome analysis reveals that *rapgef1b* plays a crucial role in the development of paraxial (somitic) mesoderm and possibly, its derivatives.

### *Rapgef1b* is required for presomitic mesoderm patterning and somitogenesis

Transcriptome analysis revealed that *rapgef1b* plays an important role in somitogenesis and mesoderm development. Hence, we examined somite development and the myogenic program in *rapgef1b* morphants at 1dpf. The phalloidin 568 staining showed disruption of the F-actin positive muscle fibres in the *rapgef1b* morphants (Fig. 5A) (*64*). To ascertain the role of *rapgef1b* in myogenesis and somite patterning, we examined the morphants for *myoD1* expression. *MyoD1* is expressed in the somites and is required for the establishment and maintenance of somite-derived muscle progenitor lineages (*65*, *66*). *Rapgef1b* morphants showed defects in the somite organization and architecture. The somites appeared enlarged and irregular with imperfect boundaries (red arrowhead, Fig.5B) as compared to the periodic and sharp somite boundaries in control (red dotted line, Fig. 5B). These somitic defects were rescued in the morphants injected with h*Rapgef1* mRNA (Fig. 5B). We also examined the patterning of somitic mesoderm. *Tbxta* is expressed in the mesoderm during early development and neuromesodermal progenitors that give rise to the spinal cord and paraxial mesoderm of the trunk and tail. At later stages, *tbxta* expression is restricted to the notochord and tailbud (*51*, *56*, *67–69*). The 1dpf morphants showed notochordal defects and expanded expression of *tbxta* in the tailbud (red arrowhead, Fig. 5C). The diffused and disrupted somitic boundaries were also observed at the early somitogenic phase (red arrowhead, Fig. 5D) coupled with the loss of segmental *myoD1* expression (red asterisk, Fig. 5E). We also observed reduced expression of *myoD1* in the PSM at the bud stage, indicating that *rapgef1b* depletion resulted in impaired presomitic mesoderm patterning (red arrow, Fig. 5E). Further, we observed expanded *tbxta* expression at the bud and 6 somite stage as well (red arrow, Fig.5F) which suggests that neuromesodermal progenitor pool is maintained in the morphants and *rapgef1b* is required to induce mesodermal differentiation.

**Figure 5:**
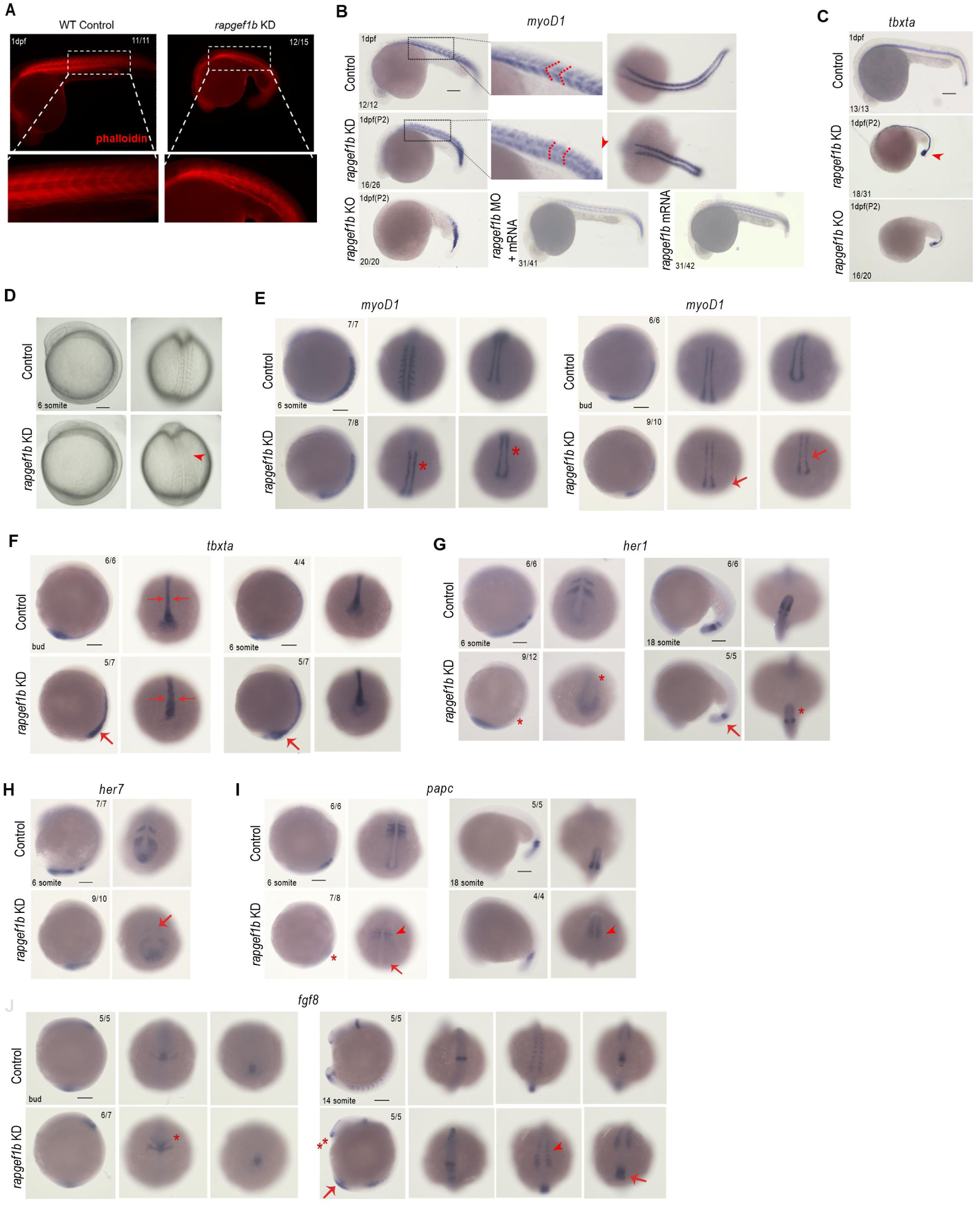
*Rapgef1b* depletion results in defective somite formation and deregulation of genes involved in presomitic mesoderm specification. **A**: Phalloidin staining in 1dpf control and morphant embryos. Inset shows the organization of F actin muscle fibers. B: *myoD1* expression at 1dpf showing somite architecture (red arrowhead, red dotted line in inset) in morphants and rescue embryos Scale bars, 200µm. C: *Tbxta* expression in the notochord and tailbud (red arrowhead) of *rapegf1b* morphants at 1dpf. Scale bars, 200µm. D: Bright field images of control and *rapgef1b* morphants at the 6 somite stage (red arrowhead). E: *myoD1* expression in control and morphants at 6 somite stage (red asterisk) and bud stage (red arrow). F: *tbxta* gene expression at bud stage (red arrow), and somite stage (red arrow). Scale bars, 200µm. G: Expression of *her1* in control and *rapgef1b* morphants in anterior somites (red asterisk) at the 6 somite stage and tail PSM (red arrow, red asterisk) at the 18 somite stage. H: Expression of *her7* in early somitogenesis in control and *rapgef1b* morphants showing reduced expression in anterior somites (red arrow). I: *Papc* expression pattern in control and morphants showing reduced expression in anterior somites (red arrowhead, red asterisk), PSM (red arrow) at the 6 somite stage, and PSM (red arrowhead) at the 18 somite stage. J: *Fgf8* expression pattern in control and morphants showing midbrain-hindbrain boundary (red asterisk) at the bud stage and in the head (double red asterisk), somites (red arrowhead), and tailbud (red arrow) at 14 somite stage.

Somites are derived from the segmentation of the PSM under the regulation of the oscillatory gene network, which involves *her* genes, Wnt, notch, and FGF signalling (*63*, *70*). *Her1* and *her7* are expressed as two stripes in the PSM, tailbud, and are essential for somite formation (*71*). *Her* deficient embryos show enlarged somites with weak boundaries. Both these genes have partial functional redundancy, yet *her1* is important for the formation of anterior somites and *her7* for posterior somites (*53*, *72*, *73*). The *rapgef1b* morphants show a lack of *her1* expression in the anterior somites in a two-stripe manner as compared to control (red asterisk, Fig.5G). At later stages, the embryos also showed reduced expression in the tail region (red arrow, Fig. 5G). We also observed reduced expression of *her7* in the somites and PSM (red arrow, Fig. 5H). Thus, the loss of segmental stripes of *her1* and *her7* suggests that the segmentation clock is disrupted in the PSM of *rapgef1b* morphants. *Papc* expressed in dorsal mesodermal cells during gastrulation contributes to the convergence movements and PSM (*74–77*). The anterior somites (red asterisk, red arrowhead, Fig. 5I) and PSM expression of *papc* is absent in the morphants (red arrow, Fig. 5I). Further, *papc* positive cells in the paraxial mesoderm are not detected in the tailbud of morphants at somite stage (red arrowhead, Fig. 5I). This downregulation of *papc* in *rapgef1b* depleted embryos suggests impaired convergence movements in the mesoderm, ultimately resulting in somitic defects. Another PSM marker, *fgf8* plays a key role in PSM maturation and somite formation (*78*). *Rapgef1b* morphants show an expansion of *fgf8* expression in the midbrain-hindbrain boundary at the bud stage (red asterisk, Fig.5J). At later stages, posterior somites appear enlarged with imperfect somitic boundaries (red arrowhead) and expanded expression in the head (double red asterisk) and tailbud (red arrow, Fig. 5J). Hence, *rapgef1b* is essential for PSM patterning, expression of segmentation clock genes and somite architecture.

### *Rapgef1b* regulates mesoderm specification by modulating canonical Wnt signaling

To investigate the underlying cause of observed paraxial mesodermal defects upregulation, we examined the Wnt/β-catenin signaling pathway in the *rapgef1b* morphants. Wnt signals are required for mesoderm induction, somite formation, and patterning through the β-catenin-Lef1/TCF pathway (*48*, *53*, *79*). The abrogation of canonical Wnt signaling impairs somitogenesis and presomitic segmentation (*48*, *50*). Indeed, during mid-blastula stage, we observed a decrease in the Wnt signaling effector β-catenin in the nucleus and cortex in interphase and mitotic phase (white asterisk, Fig. 6A). Western blotting showed a reduction in phospho GSK-3β levels (ser9) indicative of enhanced kinase activity, in turn resulting in increased phospho β catenin levels (ser33/37) in *rapgef1b* morphants (Fig.6B). Consequently, total β catenin levels decreaed in the morphants (Fig.6B), indicating the downregulation of canonical Wnt signaling. In addition, the *rapgef1b*-depleted embryos showed decreased protein levels of the Wnt effector axin2 (Fig.6B). The transcript levels of Wnt effectors *axin2*, *lef1* and *cyclin D1* were also reduced in the morphants, suggesting an overall downregulation of the canonical Wnt responses (Fig. S3F). Previous studies show that RAPGEF1 localizes to the centrosome and regulates proper centriole duplication in mammalian cells *in vitro* (*5*). Hence, we examined the role of *rapgef1b* in centrosomal dynamics during embryonic mitoses, at the mid-blastula stage. The morphants exhibited severe centrosomal and centriolar defects as shown by γ-tubulin (white asterisk, Fig.6C, S3E) and centrin 3 respectively, (Fig. S3C). Hence, *rapgef1b* is required to maintain spindle pole integrity, centrosome structure and organization during early embryonic mitoses (Fig. 6C). The *rapgef1b*-depleted embryos also showed elongated spindle length (red dotted line, Fig. 6C) and chromosome congression anomalies at metaphase (Fig. 6C). These defects are likely to result in impaired mitotic progression and chromosome instability in *rapgef1b* morphants. This is also indicated by an overall decrease in *plk1* transcript levels (Fig. S3D), essential for mitotic progression, faithful segregation of chromosomes, and proper spindle assembly (*80*). Interestingly, we observed a partial rescue of the pericentriolar defects as shown by γ-tubulin in the *rapgef1b* morphants injected with human β-catenin mRNA (white asterisk, Fig.6D). These results demonstrate that the centrosomal functions of *rapgef1b*, particularly the maintenance of spindle pole integrity are mediated by Wnt β-catenin signaling.

**Figure 6:**
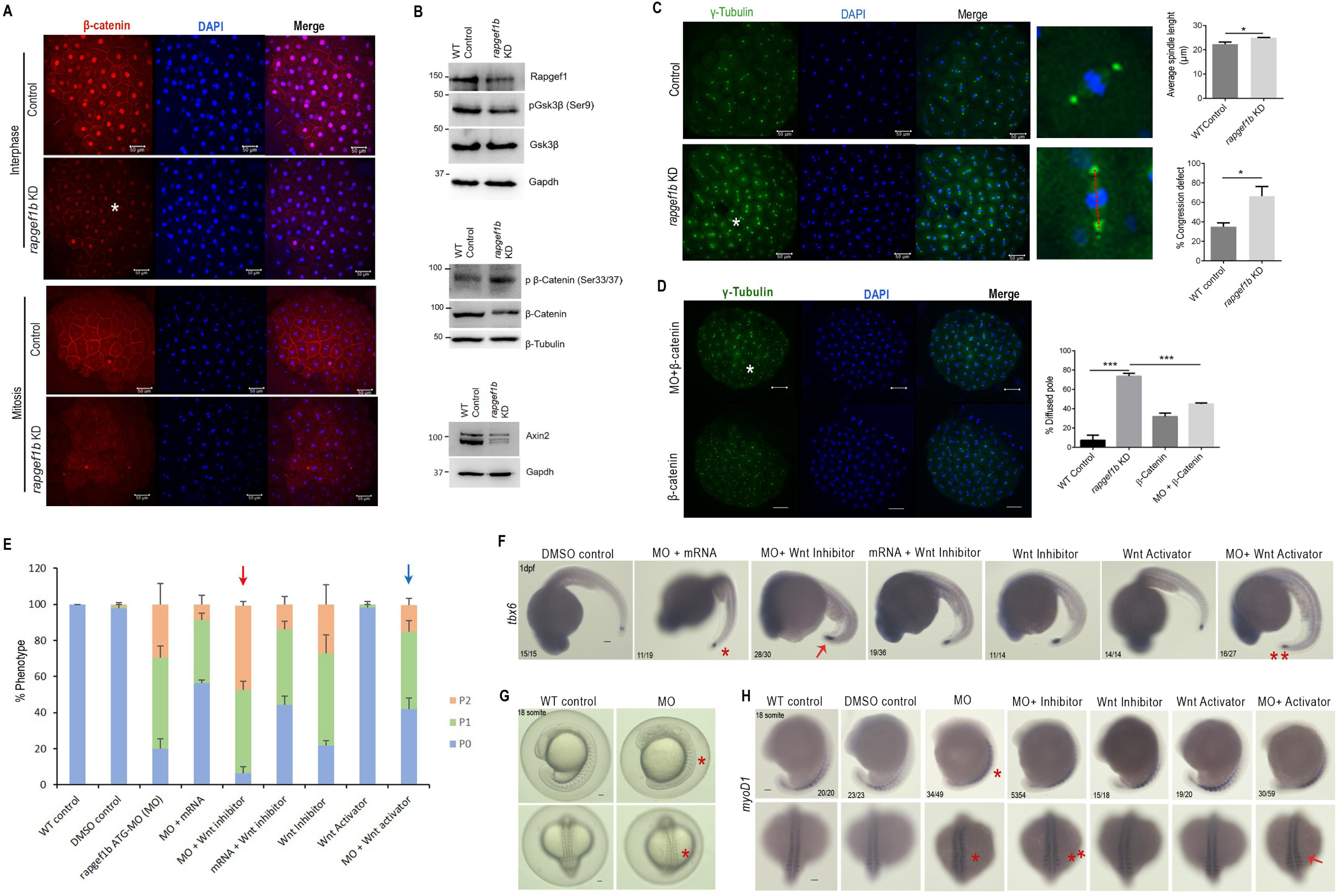
*Rapgef1b* functions by modulating Wnt/β catenin signaling during early embryonic development. A: Sum projection of confocal images of 256 cell stage control and *rapgef1b* morphants embryos showing β-catenin (white asterisk) during interphase (upper panel) and mitosis (lower panel). Scale bars, 50µm. n=3 for each experiment. Each n represents 50 embryos B. Western blotting analysis showing Rapgef1, Gsk3β/phospho Gsk3β levels, and β-catenin/phospho β-catenin levels in control and *rapgef1b* morphants. β-Tubulin and Gapdh were used as the loading controls. C: Sum projection confocal images of 256 cell stage control and *rapgef1b* morphants embryos showing centrosomal staining by γ tubulin (green) and DNA (blue/DAPI). Quantification of spindle length (red dotted line) and chromosome congression defects in the *rapgef1b* morphants as compared to control. Scale bars, 50µm. All data are shown as mean ± SEM. *p<0.05, **p<0.01, ***p<0.001. n=3 for each experiment. Each n represents 40 embryos. D: Sum projection confocal images of 256 cell stage embryos injected with *rapgef1b* morpholino and β-catenin mRNA showing centrosomal staining (white asterisk) by γ tubulin (green) and DNA (blue/DAPI). β-catenin mRNA injected embryos serve as control. Scale bars, 50µm. Quantification of spindle pole integrity (white asterisk) in the *rapgef1b* morphants as compared to control. All data are shown as mean ± SEM. *p<0.05, **p<0.01, ***p<0.001. n=3 for each experiment. Each n represents 40 embryos. E: Quantification showing percent phenotype of morphants when treated with Wnt activator and Wnt inhibitor. F: Gene expression patterns of *tbx6* (red asterisk, red arrowhead) in control, *rapgef1b* morphants, morphants treated with Wnt inhibitor and activator. WT embryos treated with Wnt activator only and inhibitor only were used as positive control. G: Bright field images of control and *rapgef1b* morphants at the 18 somite stage (red asterisk). H: Gene expression patterns of *myoD1* (red asterisk, red arrowhead) in control, *rapgef1b* morphants, morphants treated with Wnt inhibitor and activator. WT embryos treated with Wnt activator only and inhibitor only were treated as positive control. All data are shown as mean ± SEM, n=3 for each biological repeat experiment. Each n represents 20 embryos.

To further probe *rapgef1b-*mediated Wnt signaling in mesoderm patterning, we exposed morpholino-injected embryos to the Wnt activator CHIR99021 or the Wnt inhibitor XAV939 (Fig. 6E). Morphants exposed to XAV939 resulted in an increased number of P1 and P2 phenotypes (red arrow, Fig. 6E) which were rescued by administration of CHIR99021, akin to the introduction of human *rapgef1* mRNA (blue arrow, Fig. 6E). Further, expression of the paraxial mesoderm specifier *tbx6* was also restored in *rapgef1b-*depleted embryos injected with *rapgef1* mRNA (red asterisk, Fig. 6F) and CHIR99021 (double red asterisk, Fig. 6F). Conversely, *rapgef1b* depleted embryos treated with XAV939 showed an additive phenotypic expression of *tbx6* (red arrow, Fig. 6F). The somitic defects were also observed in *rapgef1b* morphants at the 18 somite stage, marking the later phase of somitogenesis (red asterisk, Fig. 6G, H). The severity of the somitic defects increased upon treatment with the Wnt inhibitor, reinforcing the synergistic action of *rapgef1* loss and attenuated Wnt signaling (double red asterisk, Fig. 6H). The somite organization and architecture were restored upon treatment with Wnt activator (red arrow, Fig. 6H). Further, inhibition of Wnt signaling also results in reduced axis elongation (*81*), which is consistent with the reduced Wnt signaling in *rapgef1b* morphants. To summarise, *rapgef1b* depletion reduces canonical Wnt/β-catenin signaling, which is the underlying basis for the impaired mesodermal patterning and somitic defects.

## Discussion

Embryogenesis is characterized by dynamic spatiotemporal gene expression that drives progenitor fate specification and subsequent differentiation. As *Rapgef1* regulates cell adhesion, cytoskeletal organization, and cell proliferation, it is well poised to play key roles in embryonic development. However, its precise role in embryogenesis remains unclear, largely because murine *Rapgef1* null embryos fail to progress beyond implantation. Hence, we investigated *rapgef1* functions in zebrafish embryos, which have two paralogs *rapgef1a* and *rapgef1b*. *Rapgef1a* depletion resulted in reduced survival rate, gross morphological defects in vasculogenesis, CNS development, and reduced locomotor activity (*13*). These zebrafish paralogs may have unique as well as overlapping functions; however, the contribution of *rapgef1b* in biological processes remains unknown.

We show that both paralogs undergo tissue- and stage-specific alternative splicing, with isoform switching evident across developmental transitions. In contrast to *rapgef1a*, the short isoform of *rapgef1b* is exclusively expressed across developmental stages, strongly indicating its unique, non-overlapping functions during early development (Fig. 1E). Further, the short *rapgef1b* isoform predominates during proliferative phases of brain development, whereas longer isoforms containing cassette exons are enriched in the adult brain, where differentiation is complete. Longer isoforms with cassette exons show differences in intra-molecular interactions which have been predicted to alter GEF activity (*19*). This temporal and tissue-specific isoform expression suggests a mechanism for fine-tuning *Rapgef1*-mediated signaling and GTPase regulation during embryogenesis. The evolutionary conservation of splicing hotspots across vertebrates further supports the functional differences between the isoforms. This also highlights some of the limitations of our current study, as isoform-specific reagents such as antibodies are not available. Future studies employing isoform-specific reagents will be critical to dissect the unique and redundant roles of these transcripts in embryonic development and to highlight their evolutionary significance. Differences in temporal expression of isoforms generated from the two paralogs may be due to the presence of alternate regulatory elements on the two linkage groups, and suggests that their products play independent roles.

Our study provides the first evidence that *rapgef1b* is essential for lineage specification in the cranial neural crest and paraxial mesoderm (Fig.7). Using CRISPR-Cas9-mediated knockout and morpholino knockdown approaches, we demonstrate that loss of *rapgef1b* disrupted early embryogenesis, neurodevelopmental and anteroposterior axis elongation defects (Fig. 2).

**Fig. 7:**
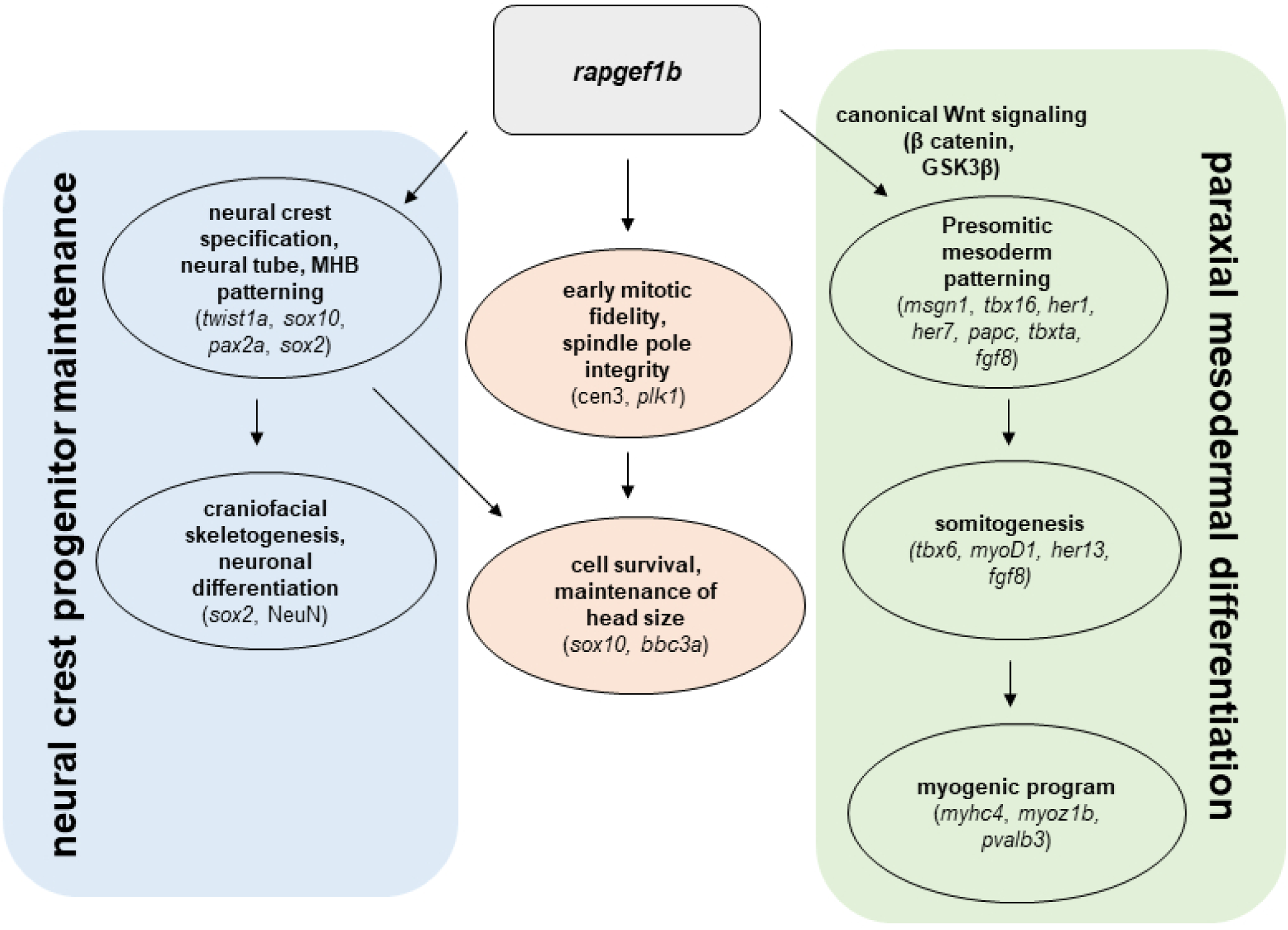
rapgef1b regulates neural crest maintenance, mitotic fidelity, and paraxial mesodermal differentiation during zebrafish embryogenesis. Schematic summary of *rapgef1b* functions in early development. On the left (blue), *rapgef1b* maintains neural crest progenitors and supports neural tube and midbrain–hindbrain boundary (MHB) patterning (*twist1a, sox10, pax2a, sox2*), which contribute to craniofacial skeletogenesis and neuronal differentiation (*sox2, NeuN*). In the centre (orange), *rapgef1b* ensures mitotic fidelity and spindle pole integrity (*cen3, plk1*), enabling cell survival and proper head size maintenance (*sox10, bbc3a*). On the right (green), *rapgef1b* modulates canonical Wnt/β-catenin signaling to regulate presomitic mesoderm (PSM) patterning (*msgn1, tbx16, her1, her7, papc, tbxta, fgf8*), somitogenesis (*tbx6, myoD1, her13, fgf8*), and myogenic differentiation (*myhc4, myoz1b, pvalb3*). Together, these findings establish *rapgef1b* as a key regulator linking progenitor maintenance, mitotic control, and mesodermal tissue differentiation.

*Rapgef1b* crispants and morphants showed reduced expression of neural crest specifier genes *twist1a* and *sox10*, increased apoptosis, and resulted in smaller head size. *Twist1a* and *sox10* mark the undifferentiated cranial neural crest and regulate the specification, maintenance of the progenitor pool, and migration (*32*, *82–84*). Further, loss of *sox10* results in apoptosis and failure to undergo differentiation to form CNS and cranial neural crest-derived craniofacial skeleton (*33*, *82*, *85*). In addition to the specification of neural crest cells, *rapgef1b* also plays a key role in the formation of neural crest derivatives. We observed the absence of neural crest-derived melanocytes in morphants and crispants (Fig. 2, 3). The transcriptomic profile also showed differential expression of genes involved in melanocyte differentiation and pigmentation (Fig. S4D). R*apgef1b* morphants showed severe craniofacial abnormalities that may be attributed to cardiac edema and the reduced expression of *sox10*, failing to undergo differentiation to form craniofacial bones and cartilage. Depletion of *rapgef1b* also resulted in mitotic abnormalities, including defective spindle pole integrity and chromosome mis-congression in zebrafish blastulae. Centriole amplification has been shown to affect the proliferation and survival of neural progenitor cells in zebrafish (*86*). These results reveal that the centrosomal functions of *rapgef1* are also conserved during embryonic mitoses in zebrafish, in a manner akin to that in mouse and human cells (*5*). Together, the reduced neural crest gene expression in *rapgef1b* morphants, coupled with enhanced apoptosis may be a consequence of impaired mitosis. The small head size, shortened interpupillary distance, mitotic aberrations, and increased apoptosis allude to a microcephaly-like phenotype in *rapgef1b* morphants. Together, these findings suggest that Rapgef1b is required to sustain progenitor identity in cranial neural crest cells and maintain mitotic fidelity during early development. Further, it is imperative to delineate the mitotic regulation by *rapgef1b* in the embryonic brain to gain mechanistic insights into its neurodevelopmental functions.

Beyond the CNS, our RNA-seq and expression analyses demonstrates that *rapgef1b* also governs paraxial mesoderm patterning, somitogenesis, and axis formation. Similar to Drosophila Rapgef1, which is required for larval muscle integrity, zebrafish *rapgef1b* morphants exhibited notochordal and somitic defects (*26*, *87*). These defects correlated with reduced expression of presomitic mesoderm (PSM) genes, particularly in anterior somites, consistent with disrupted segmentation. The PSM is derived from trunk mesodermal progenitors, and these progenitor cells are maintained in an undifferentiated state by activation of Wnt signaling and *tbxta* expression (*88*, *89*). Canonical Wnt signals are key regulators of PSM maturation and somite formation by regulating the expression of lineage determinants (*78*). Mammalian RAPGEF1 interacts with β-catenin as well as GSK3β, modulating Wnt signaling (*7*, *8*). The *rapgef1b* depletion results in the downregulation of Wnt signals, disrupting the balance between proliferation and differentiation, which induces premature pro-differentiation signals in mesodermal progenitor cells. Hence, *rapgef1b* morphants show disrupted PSM specification and defects in somite organization and architecture. Given that PSM progenitors are maintained by canonical Wnt signaling, our pharmacological experiments show that *rapgef1b* modulates this pathway, potentially via GSK3β and β-catenin regulation. *Rapgef1b* depletion dampened Wnt activity and decreased expression of Wnt effectors, leading to premature differentiation signals in mesodermal progenitors. In addition, decreased Wnt/β-catenin signaling reduced the expression of neural crest specifiers, showing that Wnt signaling coupled with BMP signals is essential for the expansion of neural crest progenitors and maintains their multipotency (*90*). The downregulation of Wnt/β-catenin signaling also resulted in a surge of apoptosis in the CNS, consistent with prior reports linking Wnt loss to cell death in the midbrain and cerebellum (*51*, *91–94*). In fact, β-catenin signaling promotes differentiation of neural crest cells into pigment cell types and suppresses neuronal cell fates in premigratory neural crest. Intriguingly, the transcriptome profiling of *rapgef1b* morphants also showed downregulation of genes involved in melanocyte differentiation and pigmentation, along with an overall decrease in neural crest-derived melanocytes in morphants and crispants. This is suggestive of *rapgef1b*-mediated attenuation of β-catenin signaling, resulting in decreased expression of melanocyte differentiation-associated genes.

In summary, our study involving detailed spatiotemporal gene expression analyses, transcriptome profiling and pharmacological intervention experiments establish *rapgef1b* as a critical regulator of cranial neural crest and mesodermal lineage specification, acting in part through modulation of Wnt signaling. The data highlight the role of *rapgef1b* in maintaining progenitor states, ensuring mitotic integrity and survival during early development. Our findings show that *rapgef1b* paralog is essential for early development, though there is the presence of a duplicate gene with high homology. Yet, we do observe some overlapping redundant phenotypic attributes such as reduced survival rate, gross morphological brain and body axis defects in *rapgef1b* crispants, as reported for *rapgef1a* morphants (*13*). Thus, a detailed analysis of the unique and redundant functions of the two paralogs of *rapgef1* in zebrafish is imperative to gain mechanistic insights into the developmental significance of *rapgef1* vasculogenesis, somitogenesis and motor neuron activity. Besides the independent functions of the two paralogs, the phenotypic effects shown by the knockdown or knockout approaches may be due to the insufficient dosage effects. Lastly, as lineage specification is also regulated by chromatin remodeling, a function attributed to RAPGEF1 in mammalian cells, it will be interesting to determine how *rapgef1b* regulates chromatin modifications to alter the expression of lineage determinants during zebrafish embryogenesis.

## Methods

### Zebrafish lines, MO injection, and characterization of phenotypes

Tubingen strain (TU-AB) zebrafish were raised according to standard protocols as described earlier. All experiments were performed according to protocols approved by the Institutional Animal Ethics Committee of the Council of Scientific and Industrial Research, Centre for Cellular and Molecular Biology, India. All the experiments were performed for a minimum of three biological replicates, and statistical analyses are stated in the figure legends. Embryos were obtained from the natural spawning of adult fish, kept at 28.5°C, and staged according to hours after fertilization (*95*). The endogenous *rapgef1* levels were depleted by using morpholino (*rapgef1b* translation blocker, 5’-CTTGCTTTCTATTTTCCCCGACATG-3′ (Gene Tools), 0.25mM, 0.5mM and 1mM in each embryo and standard control MO, 5′-CCTCTTACCTCAGTTACAATTTATA-3′ (Gene Tools), 0.25mM, 0.5mM and 1mM was used as negative controls. All the assays were performed with 1mM translation blocker unless stated otherwise. For rescue experiments, 50pg of human RAPGEF1 or β-CATENIN mRNA was co-injected along with *rapgef1b* MO in each embryo at the one-cell stage. The embryos were then analyzed for gross morphological defects and survival at later stages of development (1dpf).

### Generation of the r*apgef1b* knockout line

Single guide RNAs (sgRNAs) targeting *rapgef1b* (Ensembl ID: ENSDART00000123759.3) for CRISPR/Cas9-mediated knock-out were designed using CRISPOR online tool (https://www.crispor.tefor.net/). Two sgRNAs targeting the exons 3 and 4 of the *rapgef1b* gene were used with the following target sequences: 5′-GAGATACTTTAAGACGATTG -3′ and 5′-GCTTGGCGACTCTGATTCGC-3′. Gene-specific oligos were annealed in a PCR machine, ramping temperature down from 95°C to 4°C in a stepwise manner and run on agarose gel electrophoresis to ensure correct annealing. Both the sgRNAs were cloned into pDR274 plasmid (Addgene #42250). RNAs from both clones were *in vitro* transcribed using the Maxiscript SP6/T7 kit (AM1322) according to the manufacturer’s instructions. One-cell stage zebrafish embryos were injected with 3nl of a solution containing 600pg of Cas9 protein (TrueCut™ Cas9 Protein v2 #A36498) and 150pg of each sgRNA. The CRISPR/Cas9 approach generated a deletion of 366□bp, resulting in a premature STOP codon in exon 4. For genotyping, genomic DNA was obtained by incubating the samples (whole embryos or adult caudal fin fragments) in TE buffer supplemented with 1mM EDTA and 10 µg/ml Proteinase K (Cat no. 193981) for 1□h (embryos) or 4 h (fins) at 55□°C and 10□min at 95□°C, and then stored at 4□°C. Primers sequences used for genotyping can be found as Fig S5 (*96*).

### RNA isolation, Semi□quantitative RT□PCR, and sequencing

Total RNA was isolated from 100 embryos for each group using the RNA isolation kit (MN; 740955.50) as per the manufacturer’s protocol. Total RNA from various zebrafish adult tissues was prepared using RNAiso Plus (TaKaRa, cat. no. 9109) according to the manufacturer’s protocol. cDNA was synthesized from total RNA using PrimeScript 1^st^ strand cDNA synthesis kit (TaKaRa, cat. no 6110A). PCR was carried out using the prepared cDNA for amplification of zebrafish *rapgef1* isoforms and *rpl7* (internal control). Zebrafish isoform-specific primers for *rapgef1a* and *rapgef1b* were used to identify splice variants of these isoforms present in embryonic and adult zebrafish tissues. All primers used in this study are listed in Fig. S2. PCR was performed in BioRad C1000 Touch thermal cycler with, Taq DNA Polymerase (TaKaRa, cat. no. R500A). The following PCR conditions were used for amplification of *rapgef1*: initial denaturation was at 94 °C for 5 min, followed by 30 cycles of denaturation at 94 °C for 30 secs, annealing at 60°C for 30 secs, and extension at 72 °C for 30 sec. The final extension was for 10 min at 72 °C. Amplified PCR products were examined and pictures were captured using the Vilber-Lourmat Gel Documentation system (Germany). The PCR fragments were eluted using Macherey-Nagel NucleoSpin Gel and PCR clean-up kit (MN; 740609.50) as per the manufacturer’s protocol and sequenced to confirm that the amplicons are generated from alternate splice variants. All primers used in this study are listed in Fig.S5.

### Antisense riboprobe preparation

For cloning sequence-specific exonic fragments to synthesize riboprobes, total RNA was isolated from 1dpf embryos of the TU-AB strain using an RNA isolation kit (MN; 740955.50). The total RNA was used to prepare cDNA using the PrimeScript 1^st^ strand cDNA synthesis kit (TaKaRa, Cat. # 6110A). Sequence-specific primers were used to amplify the specific gene fragment and cloned into the pGEMT easy vector and the sequence was verified. The sequence-verified plasmid was linearised using Nco1 and antisense digoxigenin (DIG)-labeled riboprobes were synthesized using the DIG RNA labeling kit (Roche #11175025910) (*97*). All primers used in this study are listed in Fig.S5.

### Whole Mount RNA in Situ Hybridization

Embryos of various stages were fixed in 4% paraformaldehyde overnight followed by methanol fixation. Methanol-fixed embryos were rehydrated with 50% MeOH/PBT (1× PBS and 0.1% Tween 20). Embryos were rinsed with PBT, a 1:1 PBT/hybridization wash, and then a hybridization wash (50% formamide, 1.3 × SSC, 5 mM EDTA, 0.2%, Tween-20, H2O). Embryos were incubated with a hybridization mix (50% formamide, 1.3 × SSC, 5 mM EDTA, 0.2%, Tween-20, 50 μg/mL yeast t RNA, 100 μg/mL heparin, H2O) for 1 h at 55 °C. The riboprobes prepared in the hybridization mix were added and incubated for 16 h at 55 °C. Then, embryos were rinsed with prewarmed hybridization wash for 30 min (2 times) at 55 °C and followed by 1 × TBST wash (5 M NaCl, 1 M KCl, 1 M Tris, pH 7.5, and 10% Tween 20) at room temperature. Then, blocking was done with 10% FBS heat-inactivated fetal bovine serum (Gibco; 16210-064) for 1 h at room temperature. The embryos were then incubated with an Anti-DIG-AP antibody (Roche 11093274910) with a dilution of 1:5000 overnight at 4°C. The next day, embryos were transferred into a 6-well plate and washed twice with 1X NTMT (0.1M NaCl, 0.1 M Tris-Cl at pH 9.5, 0.05M MgCl2, 1% Tween-20 and H2O) for 10 min; then, incubated with 1:1 NBT-BCIP solution (Cat no. PI34042) NTMT solution for the color development. The embryos were monitored during this color reaction and stopped with 1× PBT as a stop solution. Images of embryos were captured using a Zeiss Stemi 2000CVR bright-field microscope at 5× magnification (with AxiocamICc1)(*98*) The embryos after *in situ* hybridization at 3dpf were cryoprotected with 30% sucrose in PBS, embedded in Tissue freezing medium (Leica) and cryosectioned at 15 µm thickness.

### Western Blot analysis

Whole zebrafish embryos (n=100) and adult tissues were homogenized using RIPA buffer (20mM Tris–HCl, pH 7.4; 150mM NaCl; 5mM EDTA; 1% NP-40; 0.4% sodium deoxycholate, 0.1% SDS and 1× Protease inhibitor cocktail) and kept on ice for 15min with vortexing 3 times. The homogenized lysates were centrifuged (12000 rpm for 20 min at 4°C), and supernatant were taken for the analysis. Subsequently, samples were boiled in 1× Laemmli buffer and subjected to SDS-PAGE. 20 μg of total protein was loaded in each lane on 8% SDS-PAGE gel for detection of endogenous proteins. Proteins were transferred to PVDF membranes (Merck Milipore, ISEQ00010) by electro-blotting. Membranes were blocked for 2 h at room temperature in TBST (20 mM Tris HCl, pH 7.5, 150 mM NaCl, 0.1% TWEEN 20) containing 3% BSA and probed overnight with primary antibodies (Fig. S6). After three washes with TBST, the blots were incubated with HRP-conjugated secondary antibody for 1 hr, and then washed in TBST. The blot was developed for chemiluminescence signal using the chemiluminescence (Bio-Rad) and captured in the Vilber-Lourmat Chemiluminescence System (Germany). Densitometry analysis was done using Image-J. β-Tubulin was used as a control for the normalization of protein in all the samples. The data were plotted as mean normalized protein level relative to WT± SD was plotted as error bar. Data from at least three independent experiments were plotted and significance was calculated with unpaired t-test using Prism 8.0. All the antibodies used are listed in Fig. S6.

### Real-time PCR

For quantitative real-time PCR (RT-PCR), we performed PCR using Power SYBR Green PCR Master Mix (Applied Biosystems) with ViiA 7 Real-Time PCR System (Applied Biosystems) according to the manufacturer’s instructions. Quantification data are represented as means ± SEM of three independent experiments. Analyses on normalized data were performed using the 2−δδCT algorithm (the delta−delta-Ct or ddCt algorithm). All genes were normalized against z*rpl7* unless mentioned otherwise. The GraphPad Prism software was used for the analysis. For the statistical significance of the data, p-values were calculated by performing an unpaired Student t-test. All primers used in this study are listed in Fig.S5.

### Immunohistochemistry of zebrafish embryos

Embryos of various stages were fixed in 4% paraformaldehyde overnight followed by methanol fixation. Methanol fixed embryos were rehydrated with 50% MeOH/PBST (1X PBS and 0.1% Triton X-100) followed by incubation in blocking solution 1% BSA/PBST for 2 h at room temperature. Then the embryos were incubated with primary antibody overnight at 4 °C. Next day 1X PBST washes, appropriate secondary antibody incubation overnight at 4 °C. After washing, DAPI (1µg/ml) was added for 10min followed by washing with 1X PBS and stored at -20 °C until confocal/fluorescence imaging (*98*). The mitotic phenotypes were quantified using 3D reconstruction of the confocal images using the IMARIS software. All the antibodies used are listed in Fig. S6.

### Acridine orange staining

Acridine orange (AO) staining was performed to detect the apoptotic cells in 1dpf embryos. Embryos were dechorionated, and stained with 5mg/L AO for 1 hour in the dark followed by washing with embryo water. Then the embryos were anesthetized with tricaine and were photographed by a Zeiss AxioZoom.V16 microscope, the bright green dots in the embryos indicate apoptotic cells. The number of apoptotic cells in the head region was counted using Image J software (*99*).

### Skeletal preparations by Alcian blue and Alizarin red staining

Wild-type control and morphants 5-day-old larvae were fixed in 4% PFA/PBS overnight at 4^0^C. The embryos were washed with 1XPBS and stored overnight in 100% methanol at - 20^0^C. The following staining solutions were prepared - Solution A (0.04% Alcian blue, 125mM MgCl2 in 70% ethanol) and Solution B (1.5% Alizarin Red S). 1ml of solution A and 10ul of solution B were added to the larvae and kept overnight at room temperature with constant rocking. The staining solution was removed and the larvae were washed two times with water, followed by bleach solution for 1 hour at room temperature and a gradation of Glycerol-KOH. The embryos were then stored in 50% glycerol/ 0.25% KOH for 2 hours at room temperature. The stained larvae were then imaged using a Zeiss Stemi508 stereomicroscope. For microcephaly analyses of zebrafish 5dpf larval head area was defined by the otic vesicle and the semicircle of eyes as posterior and lower boundary, respectively. The interpupillary distance (IPD) and distance between irises are also measured and quantitated using ImageJ software (*100*).

### Phalloidin Staining

Embryos were dechorinated and fixed using 4% paraformaldehyde for overnight at 4°C. Embryos were washed out of 4% paraformaldehyde at least three times with Phosphate Buffered Saline-0.1% Tween20 (PBT). Prior to phalloidin staining, embryos were permeabilized for 30min at room temperature with PBS-1%Trition. Embryos were incubated in Phalloidin–Tetramethylrhodamine B isothiocyanate (cat no.P1951) at 1∶500 dilution overnight at 4°C. Embryos were washed three times in PBS-0.1%Tw prior to imaging.

### Pharmacological modulation of the Wnt/**β**-catenin signaling pathway

Zebrafish embryos were treated at shield stage with 10 µM XAV939 and 10 µM CHIR99021 dissolved in DMSO, to inhibit (XAV939) or activate (CHIR99021) Wnt/β-catenin signalling. At 24hpf the percent phenotype was calculated for all these treated groups of embryos.

### RNAseq-based gene expression analysis

The total RNA was extracted from 1dpf control and *rapgef1b* depleted embryos using an RNA isolation kit (MN; 740955.50). Sequencing libraries were generated using MGIEasy RNA library prep set (MGI) according to the manufacturer’s instructions. Here, 500 ng total RNA was used as starting material, and rRNA was depleted by MGIEasy rRNA depletion Kit. The rRNA-depleted samples were fragmented, reverse transcribed and the second strand was synthesized and converted to cDNA. DNA was then purified using DNA Clean Beads provided in the kit followed by end repair and A-tailing. Barcoding and adaptor ligation were performed and the samples were purified. Samples were amplified using adaptor-specific primers and quantified using a Qubit dsDNA high-sensitivity kit (ThermoScientific). Sample fragment size was determined using 4200 TapeStation (Agilent). 1pMol of the dsDNA library was denatured and circularized to make single-stranded circular DNA. DNA Nano Balls (DNBs) were created using Rolling cycle amplification. DNBs were loaded onto patterned flowcell and sequenced using PE100 recipe on MGISEQ-2000 sequencer (MGI). The raw sequence data from the control and *rapgef1b-depleted* embryos were evaluated for read quality status using FASTQC (*101*). The RNAseq data was uploaded on Gene Expression Omnibus (GEO), accession no. GSE283261. Adapter trimming and read filtration were performed using CUTADAPT, and the filtration criteria were set to a minimum length (50bp), and a minimal quality score (Q20) cutoffs. The trimmed reads were then mapped onto the zebrafish fish reference genome (GRCz11) using STAR aligner (*101*). STAR was run in quant-mode, instructing the tool to generate the read counts and alignments. The differential gene expression was performed using the DESeq2 package (*102*), assayed across the three replicates of each condition; control (WT) and *rapgef1b* knockdown (*rapgef1b* KD). Transcripts with a raw count sum of less than 10 were excluded for downstream analysis. A correlation matrix was calculated to determine the agreement of various replicates across conditions. To ensure consistency across conditions and the replicates, principal component analysis (PCA) was carried out of deseq2 normalized counts. Significantly differentially expressed transcripts were obtained by filtering for a false discovery rate (FDR) by filtering p-adjusted cutoff of p <= 0.05. These are then classified as upregulated and downregulated based on their respective log fold change (L.F.C) values, which is representative of their regulation states; transcripts that had a fold change greater than 1 (L.F.C>=1) were considered as up-regulated while those lesser than -1 (F.C<=-1) were deemed down-regulated. Gene ontology (GO) enrichment analysis was performed using the enrichGO module of the ClusterProfiler package to analyze the effect of these gene subsets on metabolic networks (*103*).

## Acknowledgments

We thank Ms. Valli Undamatla from the NGS Facility, CSIR-CCMB, for help in NGS sample preparation and sequencing. We thank Dr. P. Chandra Shekar, CSIR-CCMB, for XAV939 and CHIR99021.

## Author’s contribution declaration

TP performed the morpholino and CRISPR experiments, PCR, ISH, and analyzed the data, prepared the figures, and assisted in manuscript writing. SI performed the cloning of ISH markers, ISH, immunohistochemistry, imaging, and analyzed the data. SD performed ISH, imaging, and analyzed data. RJM analyzed the transcriptome data and prepared the figure. DTS supervised the transcriptome analysis. VR and MK developed the concept. TP, VR, and MK designed the experiments and interpreted the data. MK supervised and mentored the entire study and wrote the manuscript with input from other authors.

## Availability of data and materials

Not applicable.

## Clinical trial number

Not applicable

