## supplementary_figs for "Rapgef1 paralog-mediated regulation of Wnt/β-catenin signaling orchestrates early embryo tissue patterning and morphogenesis"

**Short title**: *rapgef1b* regulates neural crest and mesoderm development

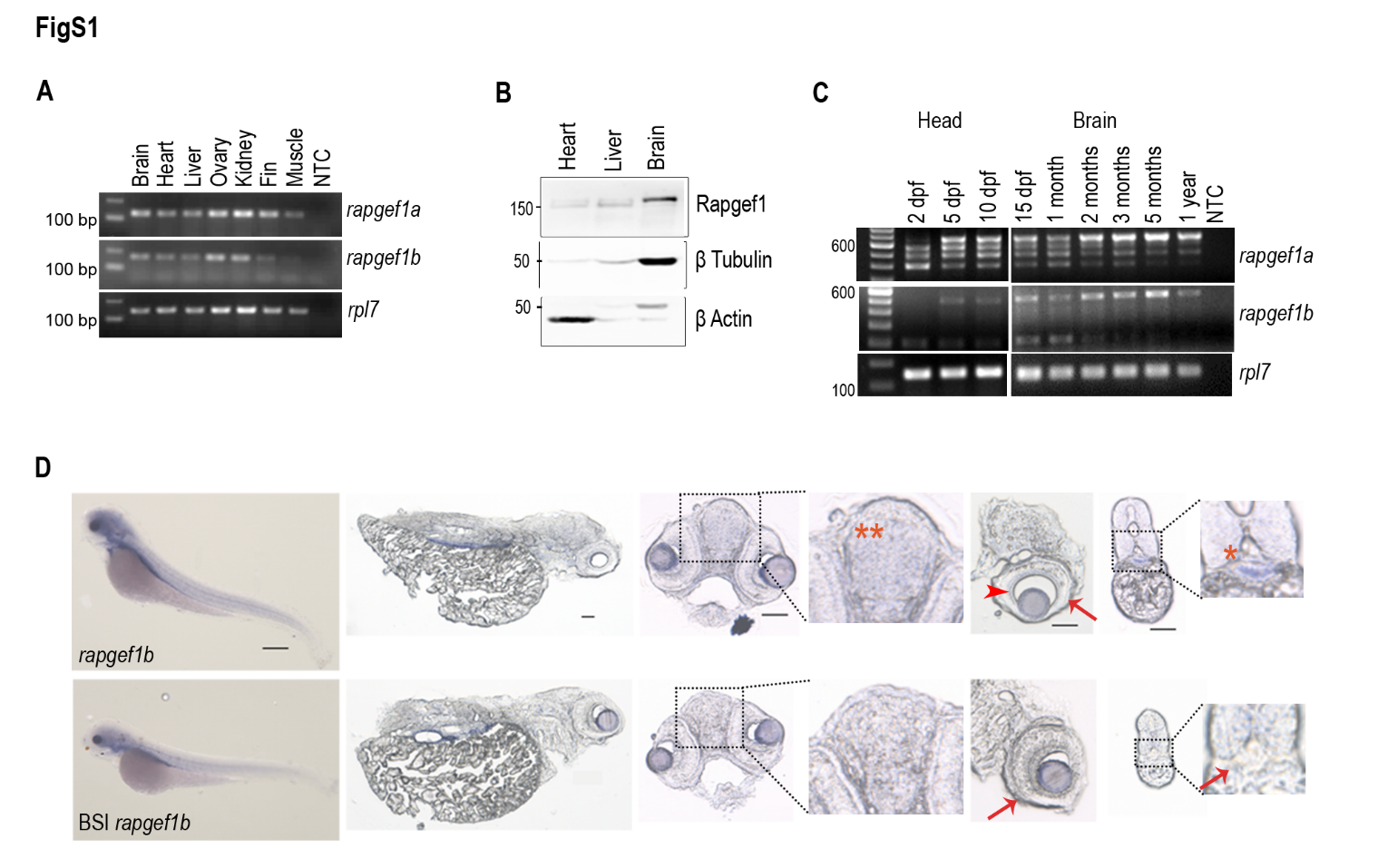

**Figure S1: Isoform-specific expression of *rapgef1* in the brain and adult zebrafish tissues**

A: RT-PCR analysis showing *rapgef1* transcripts in adult zebrafish tissues using N-ter primers for *rapgef1a* and *rapgef1b*. B: Western blot showing Rapgef1 protein levels in the heart, liver, and brain of adult zebrafish. β-Tubulin and β-Actin were used as the loading controls. C: RT-PCR analysis showing *rapgef1a* and *1b* transcripts at different larval head and juvenile stages in the brain. D: Spatial expression of *rapgef1b* and its brain-specific isoform by *in situ* hybridization and cryosections of 3dpf larvae showing diencephalon (double red asterisk), ganglion cell layer of the eye (red arrowhead), and rods and cones (red arrow). Scale bars, 50µm.

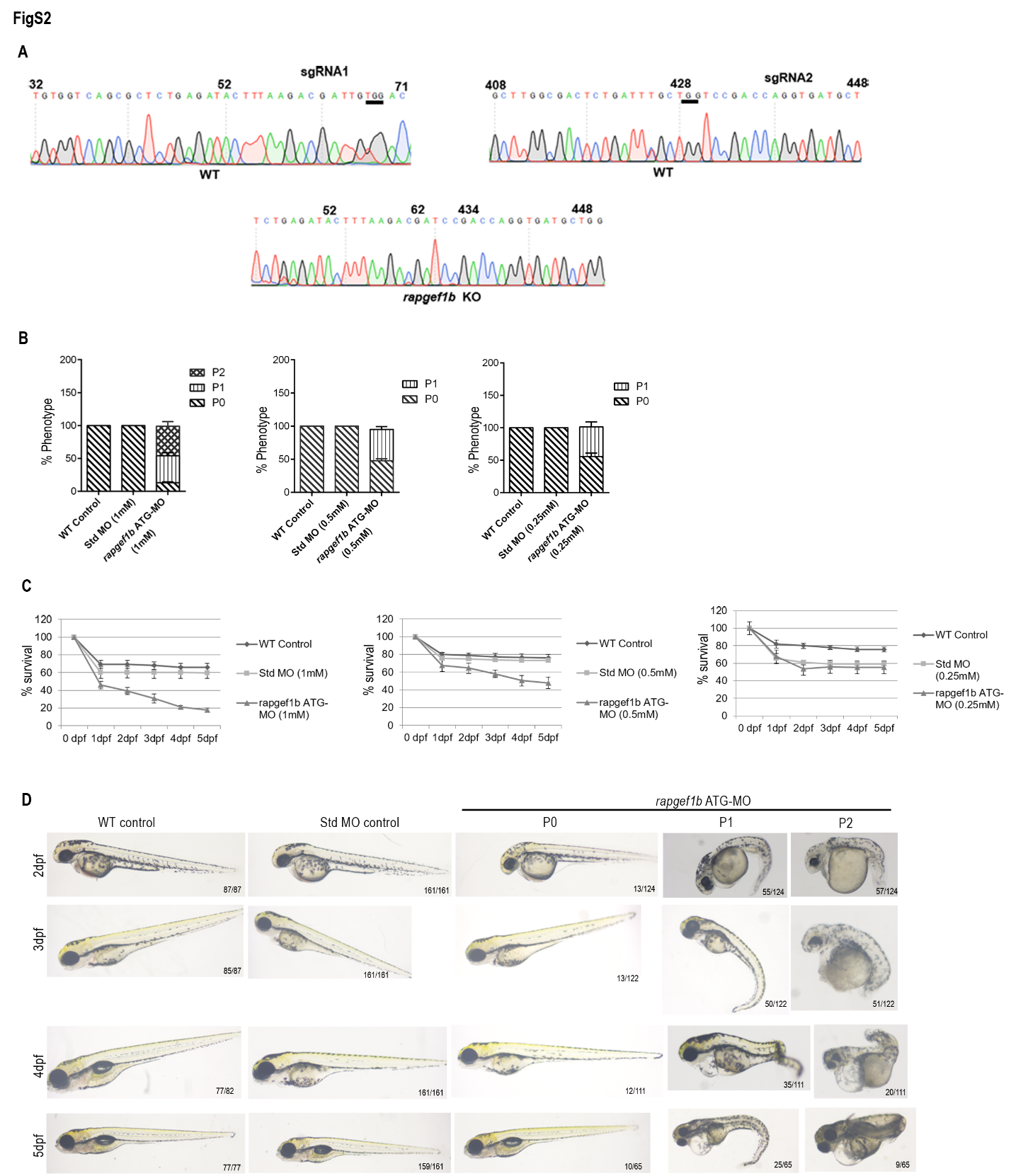

**Figure S2: Rapgef1b depletion results in developmental defects and low larval survival**

A: Snapshot of sequencing analysis highlighting the site for sgRNA in WT and *rapgef1b* knockout embryos. B: Percent phenotype of *rapgef1b* morphants in different concentrations of morpholino as compared to stage-matched control. All data are shown as mean ± SEM. n= 3 for each biological replicates. C: Percent survival of *rapgef1b* morphants over time across morpholino concentrations. All data are shown as mean ± SEM. n= 3 for each biological replicates. D: Gross morphological analysis of *rapgef1b* morphants over larval stages compared to control. n= 3 for each biological replicates and each n represents 60 embryos.

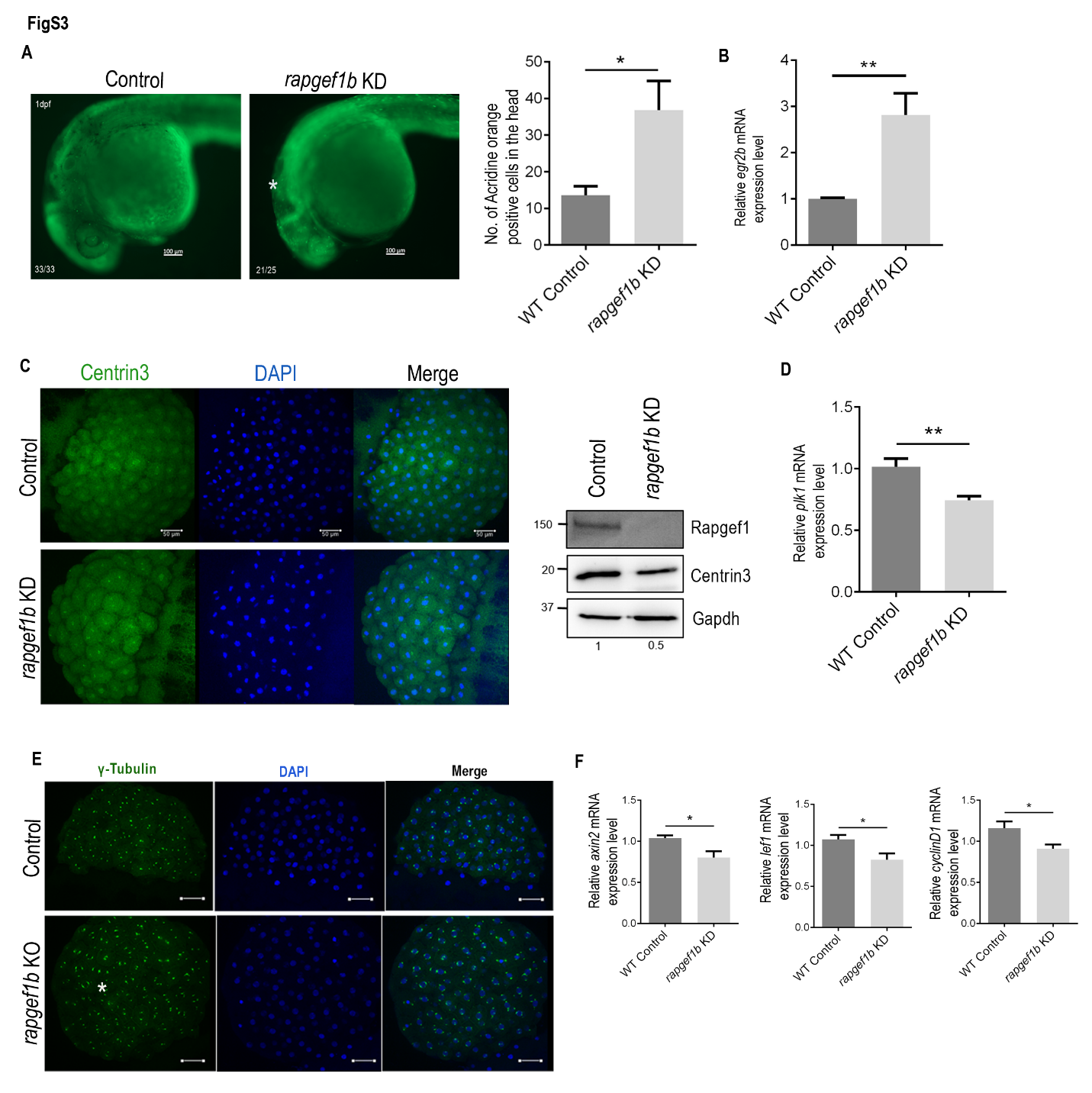

**Figure S3: Rapgef1b is essential for neural progenitor survival and spindle pole integrity**

A: Cell death analysis in the *rapgef1b* morphants by acridine orange staining. Acridine orange-positive cells (white asterisk) are shown in the head. Scale bars, 200µm. Quantitation for the number of acridine orange-positive cells in the head is shown. All data are shown as mean ± SEM. n= 3 for each biological replicate. B: Relative expression of *egr2b* in *rapgef1b* morphants. Data are shown as mean ± SEM, *p<0.05, **p<0.01, ***p<0.001. n=3 for each experiment. C: Sum projection confocal images of 256 cell stage control and *rapgef1b* morphants embryos showing DNA (DAPI, blue) and centrin3 (green). Western blot analysis to show Centrin 3 depletion in morphant embryos. Scale bars, 50µm. D: Quantification of *plk1* transcripts in control and *rapgef1b* morphants. E: Sum projection confocal images of 256 cell stage control and *rapgef1b* mutant embryos showing DNA (DAPI, blue) and γ-tubulin (green). F: Quantification of *axin2, lef1,* and *cyclinD1* transcripts in control and *rapgef1b* morphants. All data are shown as mean ± SEM. *p<0.05, **p<0.01, ***p<0.001. n=3 for each experiment. Each n represents 40 embryos.

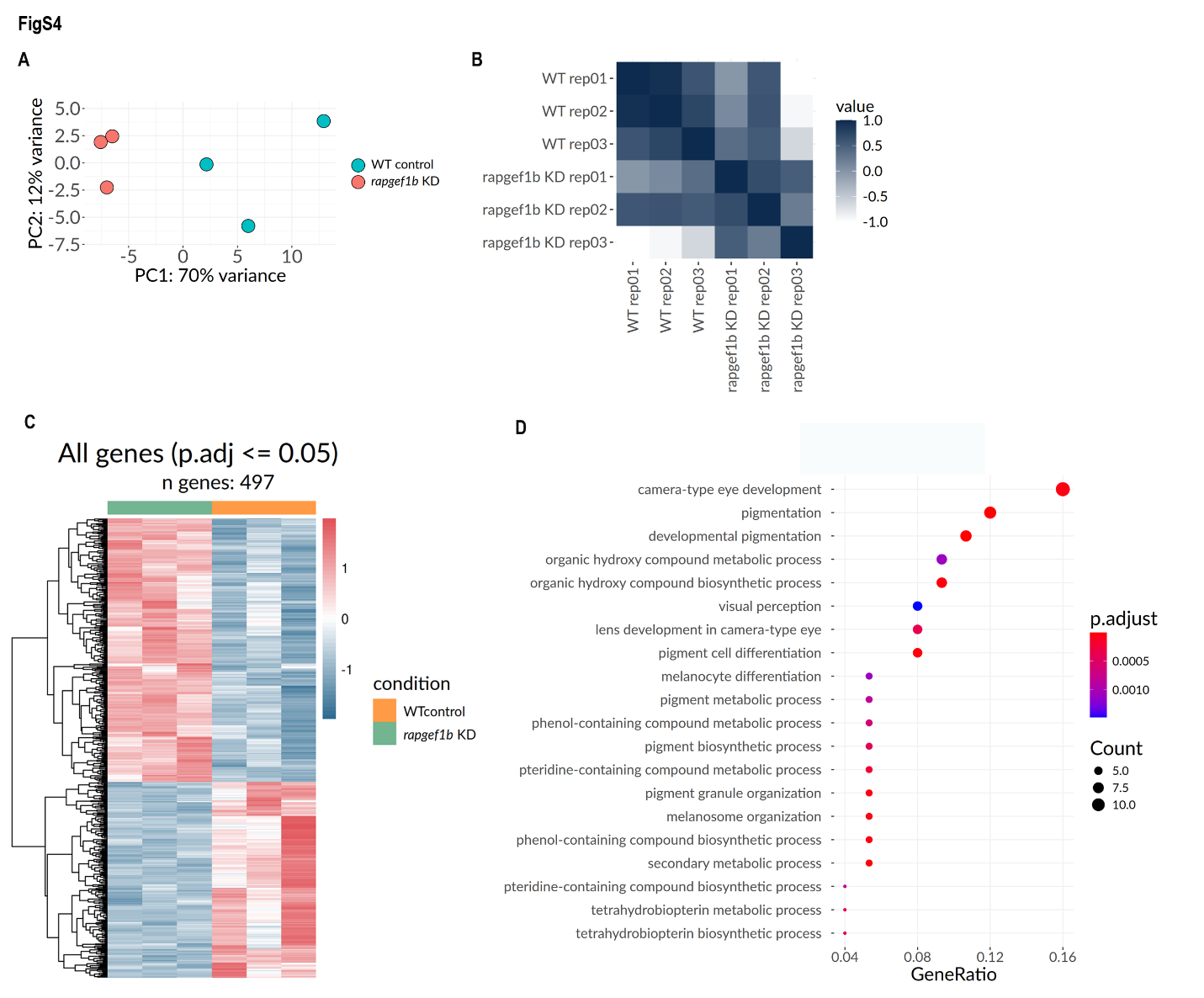

**Figure S4: Differential gene expression analysis between control and *rapgef1b* morphants**

A: Principal component analysis (PCA) RNA-seq data, PC1 represents 70% variance, while PC2 represents 12% variance. B: Pearson correlation between samples and across replicates; a correlation coefficient (R^2^) close to 1 indicates a higher relatedness, while -1 indicates unrelatedness. C: Heatmap representing differential expression patterns of 497 genes identified with a false discovery rate (FDR) of 0.05 (Q <= 0.05) across WT control and *rapgef1b* KD samples (76 upregulated and 106 downregulated genes). The colours are mapped to z-score normalized values of the log fold change across each gene set. D: Dot plot representing the gene ontology (GO) analysis of downregulated differentially expressed genes.

**Figure S5:** List of all the primers used in the study.

| **Primer name** | **Forward primer ( 5′→ 3′)** | **Reverse primer ( 5′→ 3′)** | **Purpose** |
| --- | --- | --- | --- |
| *rapgef1a*_N ter | CAAGATGGCGGTGGACAAGAA | GAGACAGGAGGACACCCTTGA | PCR |
| *rapgef1b*_N ter | GGACCCGCTGCATCAAAATG | TGAGACTTCTGGGATTCGTGC | PCR |
| *rapgef1b* | GACCCCATCTAAGAAAGG | CGTCGAGGTCGATGCCCT | ISH |
| *rapgef1a*_ISO | CGCCATATCCTGAAATACATGC | ATCAGGGCCGTCGTCGT TCT | PCR |
| *rapgef1b*_ISO | TCAACGACTCCTACAGCAGC | TGATAAAGATTCAGTGCGGTG | PCR |
| *rapgef1b*_KO | GGAGAAGGATGTGGTCAGCG | GGGGACGGTTGGTCATGTTT | KO PCR |
| *rpl7* | CTGTAGGGATAATGGCGGGTG | GTGACTTTGCTGGCCTTCTTG | PCR |
| *beta Actin* | CTCTTCCAGCCTTCCTTCCT | CACCGATCCAGACGGAGTAT | PCR |
| *twist1a* | GCCCGGTACATTGACTTCCT | TCAGGCCGAGAATCATGC | ISH |
| *sox10* | CACCACCCTCACGCTACAG | TCCACGTTACCGAAGTCGATG | ISH |
| *pax2a* | GCTGGAATGGTCCCTGGAAG | AACTTAATAACGCGGGGTTGCT | ISH |
| *sox2* | CGAGTCTAGTTCGAGTCCGC | GTTAATCGTCGTACCGGGCA | ISH |
| *tbxta* | AGACGAATGTTTCCCGTGCT | TCTGTCCTCCTCCGTTGAGT | ISH |
| *myoD1* | GCTGCCCAAAGTGGAGATTC | ATTATTCCGTGCGTCAGCATT | ISH |
| *tbx6* | CAACGCACTGATGGATGAAACC | GCGTTGAGGACACCACAGAA | qPCR |
| *cdx4* | AGTCGTCATCAACCGGCAAAA | TTCCTTAGCTCTGCGGTTCTG | qPCR |
| *cdx4* | TGGAGAAAGAGGCAAGCATGT | CTTCGTTCTCGTTTTGCCGGT | ISH |
| *tbx6* | AGTCATGTGACCCATGTGATTCC | ACAAGCAGGGATTGATTGGAAAGGT | ISH |
| *axin2* | GCTACAGGTCCTACAGACGCA | GAGTCATCCGTCAGGGCATC | qPCR |
| *lef1* | CCGCACAAGGAGCAAATCTTC | ACTGTCTAGCTGCGTCGTGA | qPCR |
| *cyclinD1* | AACTTCATCGCAAGCCCTCC | GATCGCAGACAGTCAGGGTCA | qPCR |
| *her1* | CTGCGAGAGATCAAGGCGATT | GCAGGGCACAGAGAGATACG | ISH |
| *her7* | AATCAAAATGGACAGAAAGCTGTTA | TCTGCTCGCTTGTGGTTCTT | ISH |
| *papc* | TCAGCTGAAATCCCAATTCCG | GATCAGTGGTGCCGCTTCTTT | ISH |
| *fgf8* | TTGCTACTATGCTCAGGTAACCA | GAGTAGCGGGTGCGTTTAGT | ISH |
| *pvalb1* | CTCCTTTGCTTTTCACTCGCC | TGTAGTCGAAGGAGTCGGCA | qPCR |
| *myhc4* | ACAAGCCAACTCTCACCTGTC | CCCTTCACTCTTCTTTGGCCTTTC | qPCR |
| *myoz1b* | GTGACGAGAACCTGTTGAACCT | AGCTCGTTCCCAAGGCGATA | qPCR |
| *her13* | GGATCTCAGGACATTGCTCACA | GAGTCACAGTCCCAGTTTCCTG | qPCR |
| *tbx16* | GAACAAACTCTAAAAGGCGGGC | CACCTCCAGCTCTTTACGGC | qPCR |
| *msgn1* | TGGAGGAACGTTTGCTCATCAT | AGACACCGACACTCCACTCA | qPCR |
| *egr2b* | GCGAGTGCTTCTTAGGACTTCA | TTAATCAGGCCATCTCCTGCG | qPCR |
| *bbc3* | CCGAACCATTGCCACTCAAA | CACTTCCTGTTCTGTTCCTGA | qPCR |
| *tbx16* | TGGATGAAGCAGCCCGTATC | GCTGTACACGTCTCGATGGT | ISH |
| *msgn1* | CTTCGATTCTGCCTGCTCGT | CCTAAAGCGTTGTGGGAAGGT | ISH |

**Figure S6:** List of all the antibodies used in the study.

| **Antibodies** | **Dilutions** | **Catalog number** | **Purpose** |
| --- | --- | --- | --- |
| C3G 3F6mAb |  | in-house generated (3F6) using the CBR domain of C3G | WB |
| mouse anti-β tubulin | 1:2000 | MA5-16308 | WB |
| mouse anti-β actin | 1:1000 | MA5-15739 | WB |
| mouse anti-GAPDH | 1:2000 | MA5-15738 | WB |
| anti-Mouse IgG peroxidase conjugate | 1:10 000 | A4416 | WB |
| mouse anti-α tubulin | 1:100 | DM1A mAb # 3873 | IHC |
| rabbit anti-γ tubulin | 1:2000 | PA5-34815 | IHC |
| rabbit anti-centrin3 | 1:100 | PA5-96938 | WB+IHC |
| mouse anti-β catenin | 1:200 | 13-8400 | WB+IHC |
| neuN | 1:200 | ab104224 | IHC |
| donkey anti-rabbit Alexa Fluor 488 | 1:500 | A-21206 | IHC |
| donkey anti-mouse Alexa Fluor 568 | 1:500 | A 10037 | IHC |
| sheep anti-digoxygenin-AP, Fab fragments | 1:5000 | 11093274910 | ISH |
| rabbit anti- GSK3β | 1:1000 | CST#12456 | WB |
| rabbit anti-pGSK3β (ser9) | 1:1000 | CST#5558 | WB |
| rabbit p-β-Catenin (Ser33/37) | 1:1000 | CST#2009 | WB |
| rabbit axin2 | 1:1000 | PA5-21093 | WB |
